# Confidence-guided cryo-EM map optimisation with LocScale-2.0

**DOI:** 10.1101/2025.09.11.674726

**Authors:** Alok Bharadwaj, Reinier de Bruin, Arjen J. Jakobi

## Abstract

Cryogenic sample electron microscopy (cryo-EM) maps often display uneven quality, with high-resolution features coexisting alongside weak or poorly ordered regions. Such variation complicates structural interpretation, especially for heterogeneous macromolecular assemblies. Here, we present LocScale-2.0, a context-aware map optimisation framework that operates without prior knowledge of molecular structure or composition. By leveraging general expectations of electron scattering by biological macromolecules, it enhances local detail and connectivity while preserving weak but biologically relevant structural context. We further introduce LocScale-FEM, a Bayesian-approximate deep-learning approach that emulates this optimisation to generate feature-enhanced maps. LocScale-FEM provides voxel-wise confidence scores that give a statistically grounded measure of reliability, are sensitive to local phase error in the feature-enhanced map, and highlight regions where interpretation warrants caution. Selected examples illustrate how such maps can aid biological interpretation and how confidence-guided map optimisation can increase objectivity in cryo-EM density analysis.

## Introduction

Over the past decade, cryogenic electron microscopy (cryo-EM) has transformed structural biology by enabling high-resolution structure determination of macromolecular assemblies that were previously difficult or impossible to study by other structural methods. Cryo-EM now routinely delivers near–atomic resolution reconstructions across a wide range of systems and has become a cornerstone technique for exploring the structure if molecular machines in their functional states and native environments ^1,2^. Current applications increasingly target systems that are structurally and compositionally complex: membrane proteins in native or near-native lipid environments ^3–5^, large assemblies with flexible or low-occupancy subunits ^6–8^, samples purified from endogenous sources ^9–11^ and *in-cell* structures obtained by subtomogram averaging from native cellular samples ^8,12,13^. Such specimens often contain extensive structural context such as lipid/detergent shells or embedding membranes, heterogeneous modifications and peripheral scaffolds, and exhibit pronounced conformational and compositional heterogeneity. These features are frequently weak or poorly ordered, yet often biologically important. Interpreting such reconstructions therefore requires methods that enhance high-resolution detail without suppressing weak but informative low-resolution context, and ideally provide an estimate of confidence in regions where the map is ambiguous. Automating these processes is essential for reproducibility and for making reliable interpretation accessible beyond expert users.

In cryo-EM single-particle analysis and subtomogram averaging, three-dimensional electric potential maps are assembled by averaging information of many molecules in differing orientation from low signal-to-noise ratio projection images or tomographic subvolumes. Because the ensemble of imaged molecules inevitably contains molecules in multiple states, the final reconstructions represent averages over conformers and compositions, even after classification into structurally uniform subsets. Regions of structural variability therefore exhibit reduced signal-to-noise ratio (SNR) and lower effective resolution ^14–16^. The result is, sometimes strong, spatial variation in map quality across the reconstructed volume. To compensate for the variation in resolution-dependent contrast loss, and to facilitate structural interpretation, maps are typically post-processed after reconstruction by local spectral reweighting of Fourier terms ^17–21^ and real-space filtering ^22–25^. More recently, density-modification approaches have extended these strategies by incorporating physical and statistical priors about expected map properties into post-processing ^21,26,27^ and during map reconstruction ^28,29^. We refer to this spectrum of procedures collectively as map optimisation.

Deep learning methods offer alternatives that mimic the effect of local sharpening or density-modification and produce optimised maps with little or no manual intervention ^30–32^. While these methods are widely adopted, their lack of explicit reliability estimates introduces important caveats for interpretation. Deterministic neural networks risk introducing bias by suppressing or hallucinating signal and enhancing artefacts without signalling their uncertainty, leaving practitioners to subjective judgement when deciding which map features to interpret, especially in ambiguous, low contrast regions. With the ongoing exponential growth in cryo-EM structures this poses a risk for misinterpretation, especially by new and inexperienced users ^33^. A framework that integrates map optimisation with robust confidence measures would therefore represent a critical step toward more objective and reproducible cryo-EM map analysis.

Here we introduce LocScale-2.0, an automated framework for context-aware and confidence-guided map optimisation. The method extends the original LocScale approach ^18^ with a procedure for locally adaptive map optimisation that requires no prior model information. In addition, we develop a machine learning strategy that emulates this optimisation and provides a voxel-wise confidence metric to assess the reliability of individual features. Together, these components yield maps that preserve weak but biologically informative context, improve contrast for high-resolution detail, and support more objective structural interpretation.

## Results

### Multimodal cryo-EM map optimisation in LocScale-2.0

LocScale-2.0 integrates physics-informed and deep learning-based map optimisation into complementary workflows (Figure 1). The first workflow couples model-independent local scale estimation with windowed Fourier-amplitude scaling, which is conceptually related to local map sharpening in the original LocScale approach ^18^. Instead of using explicit model information as a reference, it employs physical priors to estimate and locally compensate for resolution-dependent contrast loss. These priors draw on established expectations for the averaged squared structure factor of biological macro-molecules ^21,34,35^. The second workflow implements a deep convolutional neural network, MC-EMmerNet, that learns from pairs of unmodified input maps and maps optimised with our physics-informed procedure to predict *feature-enhanced maps* directly in real space. Unlike the physics-based method, which adjusts only the amplitudes of Fourier coefficients and leaves their phases locally unchanged, this procedure modifies both amplitudes and phases in a manner similar to density modification. To quantify predictive uncertainty, we adopt a Bayesian approximation using variational inference with Monte Carlo dropout ^36^. In addition, the workflow produces a baseline map derived from local amplitude scaling of the unmodified half maps with the mean predicted map, which serves as a reference for computing a voxel-wise confidence metric that is sensitive to local phase discrepancies and hence putative bias in the feature-enhanced map (see Methods).

**Fig. 1.**
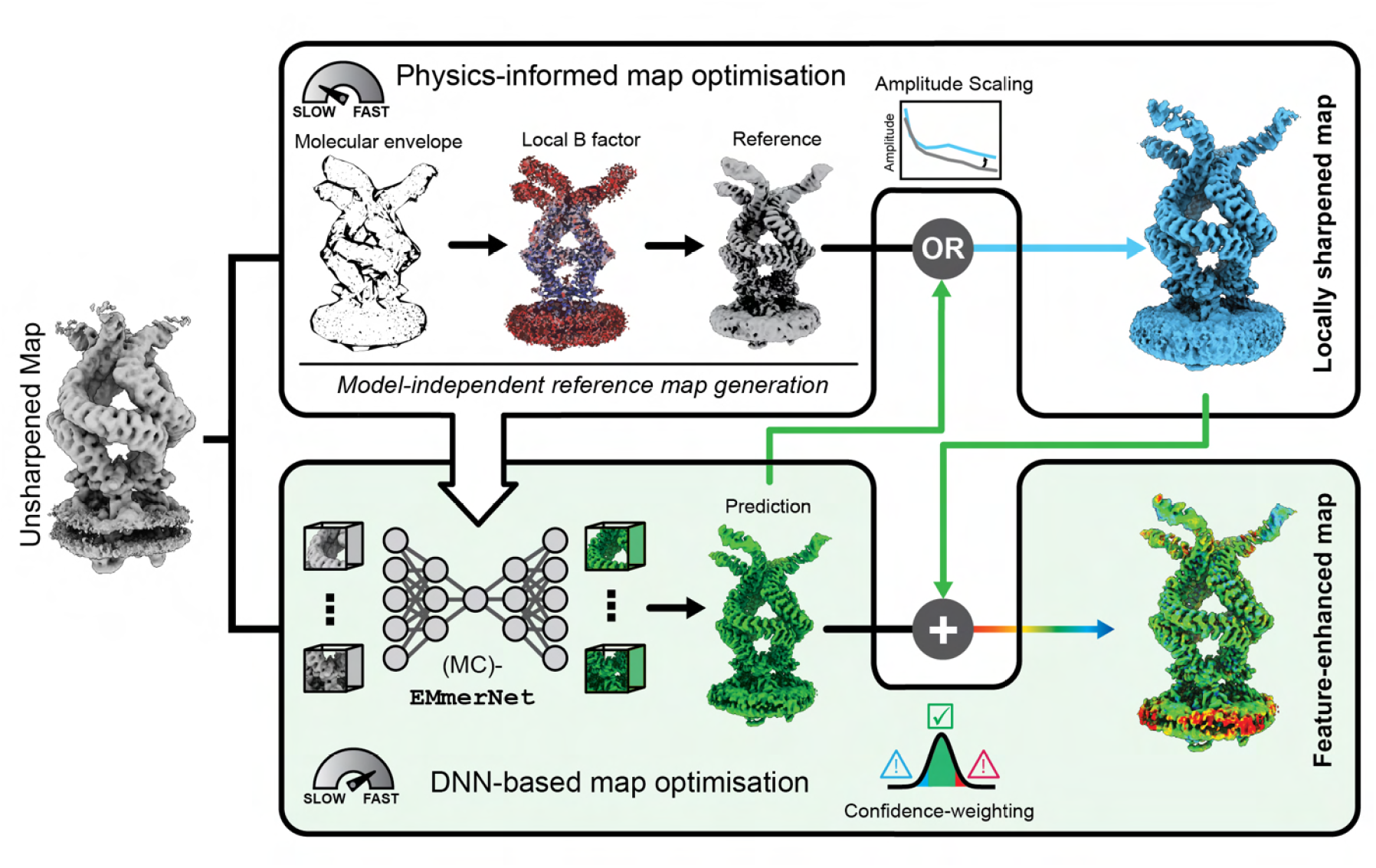
Overview of LocScale-2.0. LocScale-2.0 provides two complementary workflows for map optimisation. The physics-informed workflow leverages priors on electron scattering properties to generate reference maps, which are then used for local scaling of radially averaged Fourier amplitudes. The deep neural network (DNN)-based workflow employs MC-EMmerNet, a network trained to emulate the physics-informed process, to directly predict an optimised map. This predicted map can either serve as a scaling reference or, when generated with a variational inference-enhanced network (MC-EMmerNet), as a directly interpretable *feature-enhanced map*. Confidence scores are calibrated against amplitude-modified baseline maps, enabling systematic identification of high-versus low-confidence regions for structural interpretation (see LocScale-FEM).

### Physics-informed map optimisation

A limitation of most existing map-optimisation methods is their susceptibility to bias. This limitation affects reference-based sharpening approaches implemented in LocScale-1.0^18^ or phenix.autosharpen ^19^, as well as density modification ^27^ and deep learning-based methods ^30–32^. In reference-based methods, inaccuracies or incompleteness in the input structural model may cause suppression of density in regions not supported by the reference, and obscure relevant contextual features for biological interpretation ^18,21^. Deep learning-based methods ^30–33^ face analogous challenges, often compounded by limitations in representa-tiveness of training data and generalisability. An additional concern for these latter methods is positive bias, where plausible but non-existent structural details are hallucinated in a way that could lead to misinterpretation.

To mitigate these issues, we developed a method for local map optimisation that considers the unbiased molecular signal of the 3D reconstruction and requires no prior structural information. The approach constrains the radially averaged Fourier amplitudes using first-principles expectations and data-driven estimates of the average squared structure factor of biological macromolecules. The spectral profile of this radial structure factor depends on molecular mass distribution, local *B* factors, and secondary structure ^21,34,35^. To approximate these parameters, we first estimate the molecular volume using statistical hypothesis testing with false discovery rate (FDR) control ^37^ and define density consistent with genuine molecular signal at the 1% FDR threshold (Supplementary Figure S1a,b). Molecular mass estimates from the volume enclosed by the binarised FDR mask correlate strongly with masses computed from primary sequences and support our chosen threshold as a suitable proxy for the molecular envelope (Supplementary Figure S1c,d). Importantly, this method reliably generates envelopes that capture density from molecular components absent from the primary sequence, enabling its application to complexes with heterogeneous or unknown composition (Supplementary Figure S1c,d).

Within this envelope, pseudoatoms are placed randomly to initialise a pseudomodel, which is then iteratively refined against the experimental density (Supplementary Figure S2a–c). In many practical cases, partial structural information is available from prior experimental models or structure prediction and can be exploited in a hybrid workflow that uses explicit model coordinates where available and a pseudoatomic representation in unmodelled regions (Figure 2a–c; Supplementary Figure S1a). This strategy ensures that all density within the molecular boundary is represented and prevents signal suppression in regions lacking atomic models.

**Fig. 2.**
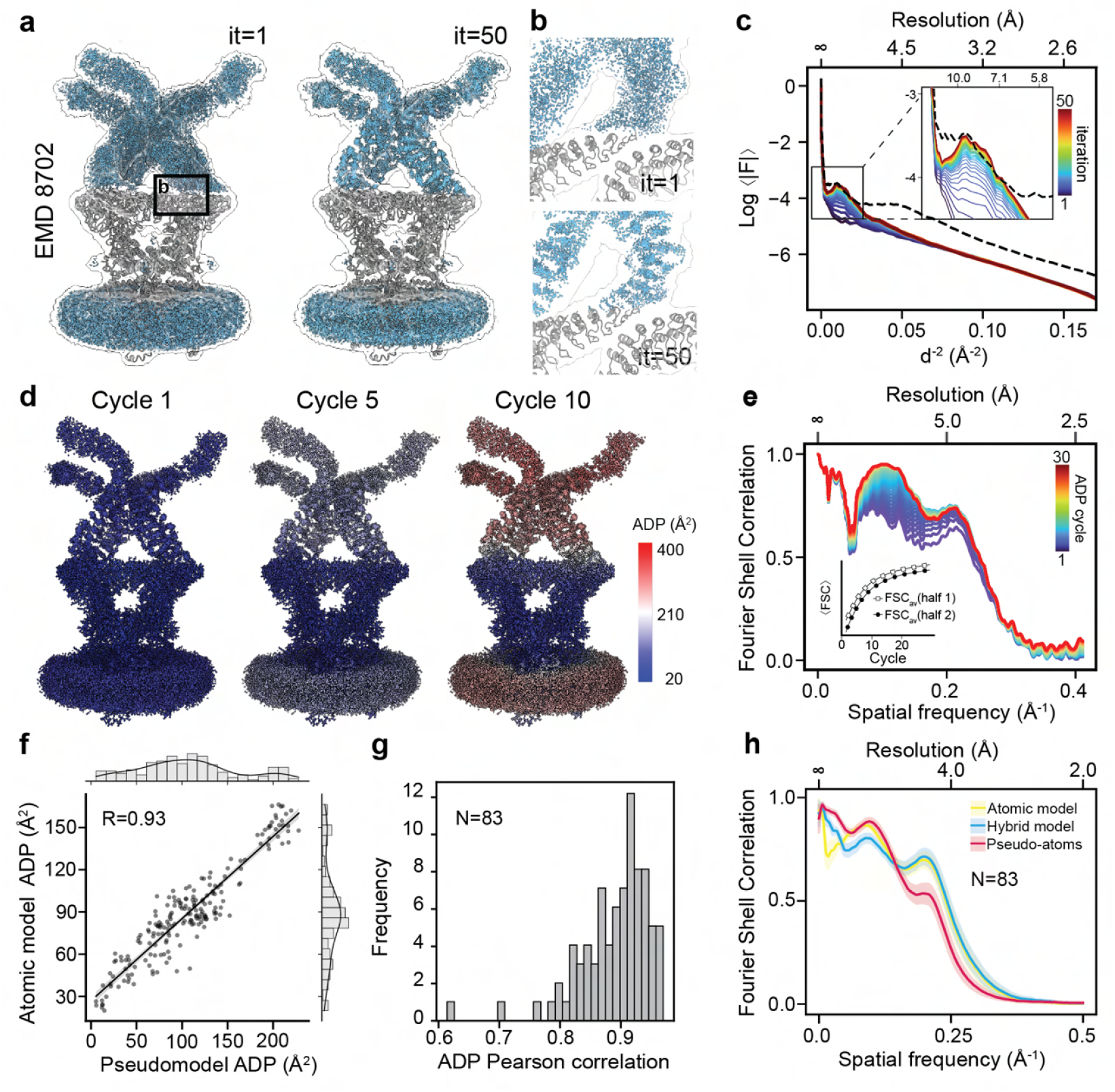
Pseudomodel generation and ADP refinement. (a) Example for pseudoatom placement and refinement for *Drosophila melanogaster* NOMPC (EMDB: EMD-8702) with partial prior model information (hybrid model). Pseudoatoms (blue) are uniformly placed within the molecular boundary defined by the FDR confidence mask except for regions for which prior model information is available (gray). Models are shown for the start (iteration 1) and end (iteration 50) of the refinement. (b) Close-up of a region at the interface between explicit and pseudomodel. (c) Radially averaged Fourier amplitudes of model maps computed throughout the iterations of pseudomodel refinement. Inset: Close-up of low-to-medium resolution region defining the overall molecular envelope. (d) Atomic displacement factors (ADP) mapped onto atomic coordinates shown for different snapshots across the ADP refinement cycles. (e) Evolution of model-to-map Fourier shell correlation (FSC) shown for a series of ADP refinement cycles. The insets shows the half-map validation displaying the average integrated FSC for work and test maps as a function of refinement cycle. (f) Correlation of atomic model and pseudomodel ADPs averaged in 25 Å windows. R denotes the Pearson correlation coefficient. (g) Distribution of Pearson correlation coefficients for window-averaged ADPs between atomic and pseudomodel representations across 83 EMDB entries. (h) Average model-to-map Fourier Shell Correlation computed for 83 EMDB entries and their corresponding PDB-deposited models, pseudomodels and hybrid models. The shaded areas correspond to 95% confidence intervals. Maps were resampled to uniform pixel size for comparison. See also **Supplementary Figure** S1-S3.

To approximate the physical decay of the overall structure factor, atomic displacement factors (ADPs or *B* factors) are then refined using restrained refinement based on local neighbourhood averaging across an empirically determined radius (Figure 2d–e, Supplementary Figure S2d–e; Methods). These restraints ensure that *B* factors of pseudoatoms, which are independent and non-bonded, vary smoothly, thereby effectively reducing the number of free refinement parameters. Refined local *B* factors from pseudomodels agree closely with those from restrained ADP refinement of atomic models (Figure 2f–g, Supplementary Figure S3d–e), without evidence of overfitting (Figure 2e, Supplementary Figure S2e). The resulting *B* factor distributions follow the shifted inverse-gamma behaviour characteristic for macromolecular structures ^38,39^. In several cases, we observed multimodal *B* factor distributions consistent with structural heterogeneity such as flexible loops, peripheral or low-occupancy subunits, detergent micelles, and with prior reports that local mixtures of shifted inverse-gamma distributions reflect conformational or compositional variability ^39^ (Supplementary Figure S3h-i). Together, these steps provide model-independent estimates of local structure factors and their resolution-dependent falloff, from which radially averaged structure factors can be computed. Because the pseudomodel representations only approximate secondary structure, we apply empirical priors to imprint expected spectral texture of secondary structure elements from proteins and nucleic acids into the local radial structure factor over the relevant frequencies ^21,29^. The spectral profiles of the locally averaged Fourier amplitudes are then used for local map sharpening as described previously ^18^. A detailed account for the implementation of the above procedure is given in the Methods.

Comparison of model–to–map Fourier Shell Correlation (FSC) curves for 83 model–map pairs illustrates the respective strengths and limitations of atomic, pseudoatomic, and hybrid reference models for local amplitude scaling (Figure 3h). Pseudomodels, which lack explicit stereochemical information, are well suited to capturing low-resolution scattering behaviour by representing signal-containing regions not typically covered by atomic models, but they cannot fully reproduce high-resolution features. Atomic models, in contrast, provide an accurate description of high-resolution signal but tend to suppress density in weak, unmodelled or heterogeneous regions. Hybrid models combine these complementary advantages and yield the most balanced representation of the experimental density. In the following we will therefore focus our discussion on applications of this hybrid approach for reference generation and subsequent local sharpening in LocScale-2.0.

**Fig. 3.**
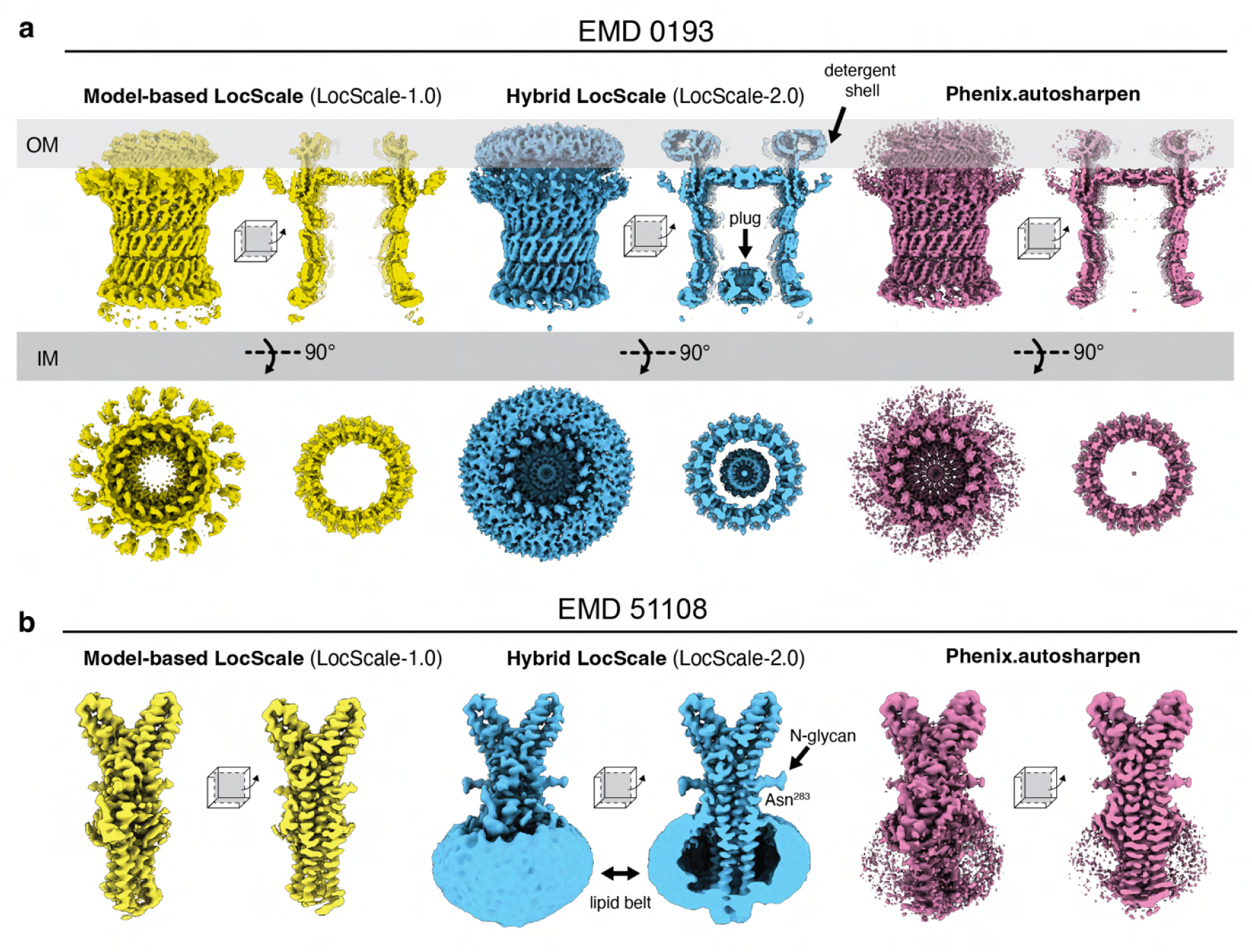
Comparison of LocScale-2.0 hybrid mode map optimisation with other local sharpening methods. (a) Map optimisation of *Klebsiella pneumoniae* type II secretion system outer membrane complex (EMDB: EMD-0193) using model-based sharpening with LocScale-1.0 (left), hybrid reference sharpening in LocScale-2.0 (middle) and phenix.autosharpen (right). Approximate locations of the outer (OM) and inner (IM) membrane are indicated. Arrows mark the visible detergent shell of the outer membrane-spanning segment and the section views reveal the central disordered plug occluding the lumen. (b) Optimised maps for human Tweety homologue 2 (EMDB: EMD-51108). The arrows mark density for N-linked glycans and the lipid belt around the the hydrophobic cavity. See also **Supplementary Figure** S4.

### Preserving structural context with hybrid map optimisation

To benchmark our procedure, we applied LocScale-2.0 to generate optimised maps for a set of representative single-particle reconstructions and subtomogram averages from the Electron Microscopy Data Bank (EMDB), and compared them with corresponding maps produced by two established local sharpening approaches: LocScale-1.0^18^ and phenix.autosharpen ^19^ (see Methods for details). Across test cases, LocScale-2.0 produced maps whose high-resolution detail was generally comparable to, and in some cases improved relative to, the alternative methods, while additionally preserving and enhancing weak but informative density in regions that are flexible, disordered, or generally unmodelled such as detergent shells, lipid belts, N-linked glycans, and low-occupancy subunits (Figure 3, Supplementary Figure S4). A notable feature of the optimised maps is that both high-resolution detail and weak, low-resolution contextual density remain visible at a single density threshold, allowing co-visualisation of these densities during map interpretatioo and facilitating direct comparison between structurally heterogeneous regions.

For example, in the *Klebsiella pneumoniae* type II secretin (EMD-0193), the LocScale-2.0 map co-visualises the plug density that occludes the lumen, a feature likely unresolved at high resolution due to its mismatch with the *C*15 symmetry of the secretin complex ^40^. In the deposited half maps this feature is only detectable after strong low-pass filtering (Figure 3a, Supplementary Figure S4e). In subtomogram averages of *Mycoplasma pneumoniae* 70S ribosomes (EMD-13234) from *in-cell* to-mograms, the LocScale-2.0 map co-visualises heterogeneous density at the mRNA entry and A-site tRNA without loss of high-resolution detail in well-resolved regions (Extended Figure S4a-c). Similarly, in Tweety homologue 2 (TTYH2; EMD-51108), LocScale-2.0 captures the largely continuous lipid/detergent belt surrounding the transmembrane region as well as the N-linked glycans, while retaining fine detail of the transmembrane helices, including individually resolved, ordered lipids (Figure 3b, Supplementary Figure S4d). Overall, these observations show that hybrid reference-based sharpening simultaneously enhances weak contextual features and high-resolution detail, and yields maps that represent heterogeneous complexes and assemblies in a manner more consistent with their structural context. This extends to highly heterogeneous assemblies: In apolipoprotein B-100 (apoB-100)-assembled low-density lipoprotein (LDL) particles (EMD-44469), for instance, the method emphasises well-resolved density for the structured apoB-100 N-terminal domain including morphological features of its β-belt formed by amphipathic β-sheets wrapped around the surface of the core lipids of LDL, while retaining the weak density consistent with cholesteryl ester plates within the LDL core (Figure 4a,b, Supplementary Figure S4f). By contrast, these weak and low complexity densities are entirely suppressed by the alternative methods.

**Fig. 4.**
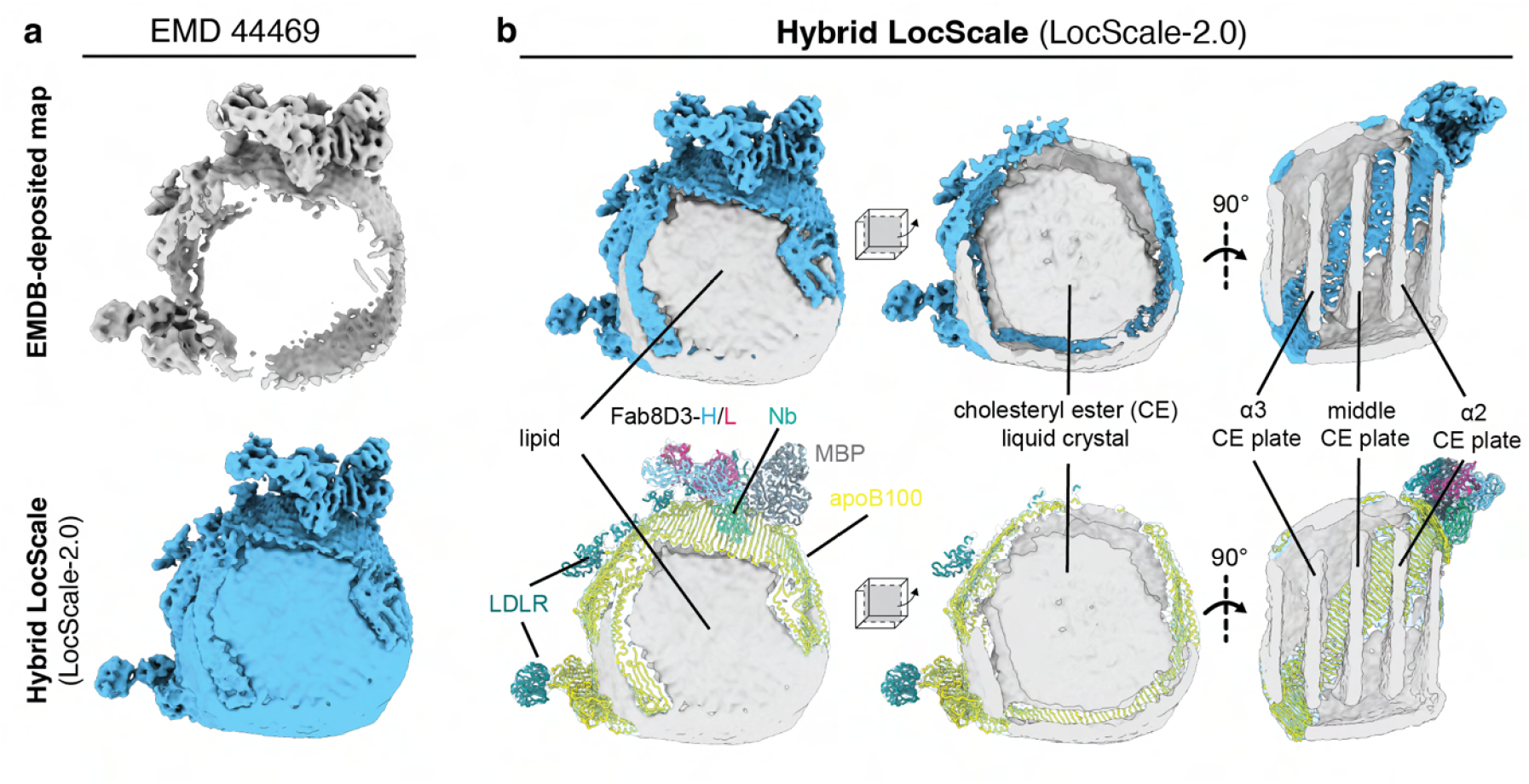
Preservation of contextual structure by LocScale-2.0. (a) cryo-EM density maps of Apolipoprotein B 100 bound to LDL receptor and legobody (EMDB: [EMD-44469](https://www.ebi.ac.uk/emdb/EMD-13234) – top; deposited ^6^) and after hybrid reference sharpening in LocScale-2.0 (bottom). (b) LocScale-2.0 optimised map EMD-44469. Top: Density assigned to protein is shown in blue; unassigned contextual lipid density is shown in grey. Bottom: The deposited atomic model (PDB: 9bdt) fitted into the optimised map with colours as labelled. Cross-sections show the cholesteryl ester plates in the interior of the low-density lipoprotein (LDL) particle. See also **Supplementary Figure** S4.

### LocScale-2.0 optimised maps aid automated model building

We next asked whether the improved balance in representing high- and lower resolution features in LocScale-2.0 optimised maps can aid automated methods for atomic model building. To test this, we used the cryo-EM model building software ModelAngelo ^41^ on a benchmark set of 50 map–model pairs from the EMDB and PDB, each determined at better than 4.0 Å FSC resolution. For each case, an optimised map was generated using the hybrid LocScale-2.0 procedure, and ModelAngelo was run both on these maps and on the corresponding EMDB-deposited maps (Supplementary Figure S5). Overall, ModelAngelo achieved higher sequence coverage from LocScale-2.0 maps than from deposited maps, both before (82%) and after (72%) pruning of short peptide fragments (Supplementary Figure S5a-c). On average, these models also exhibited smaller predicted backbone root-mean-squared deviation (r.m.s.d.) for both protein and nucleic acid components. Interestingly, improvements were observed for targets both represented and not represented in the ModelAngelo training dataset. In several cases, the improvements were substantial. For example, in the *Azospirillum brasilense* glutamate synthase (EMD-10106), the LocScale-2.0 map enabled building of 1306 additional residues compared to the deposited map (Supplementary FigureS5d,e). Given that this *α*_6_*β*_4_ complex comprises 11108 residues in total, this represents an increase of ~ 12% in modelled sequence. In 17% of the benchmark cases, however, the EMDB-deposited maps produced higher sequence coverage than LocScale-2.0 maps and in 11% the sequence coverage was equal, indicating that improvements are not uniform across the dataset. The sequence recall (i.e. the fraction of residues correctly identified compared to the PDB deposited model) computed from the 50 test map-model pairs was comparable for EMDB-deposited and LocScale-2.0 maps (Extended Figure S5f-j). Since ModelAngelo was trained on generally *B*-factor–augmented maps, these results suggest that additional gains might be realised by retraining ModelAngelo, or related deep learning–based model building tools, directly on LocScale-2.0 maps, thereby potentially extending their reach to lower resolution. Complementary to retraining, LocScale-2.0 optimisation could also be applied at the inference step, so that model building benefits from context-aware maps during prediction. In practice, these strategies may be combined to maximise performance.

### Feature-enhanced maps from uncertainty-aware deep learning

We next investigated whether the physics-informed map optimisation procedure can be replicated and accelerated using machine learning. To this end, we trained a 3D U-Net on pairs of unmodified half maps and hybrid LocScale-2.0 maps to predict optimised maps directly from unmodified input densities. Predictions from this network, which we term MC-EMmerNet, emulate the sharpening achieved by the physics-informed approach (Supplementary Figure S6a-d) at substantially reduced computational cost. For example, for a large and context-rich structure such at the apolipoprotein B100 assemblies (EMD-44469; Figure 4 complex, network prediction speeds up map optimisation about 16-fold compared to the physics-informed procedure (14 min vs. ~ 4h; Supplementary Table S1). However, deterministic networks provide point estimates with no measure of predictive uncertainty that would allow identification of regions where the output of the network may be unreliable. To overcome this limitation, we implemented variational inference in the 3D U-Net using Monte Carlo dropout, which retains dropout layers at inference time and samples multiple stochastic forward passes ^36^. This produces an ensemble of predictions from which voxel-wise means and variances can be computed, yielding both the average predicted map (which we call <u>F</u>eature-<u>E</u>nhanced <u>M</u>ap, or LocScale-FEM) and a spatially resolved estimate of predictive uncertainty (Figure 5a, Figure S6e-g). Rather than using the raw voxel-wise variance from these predictions as the uncertainty metric, we take the ensemble mean as a reference to perform local amplitude scaling and generate a baseline map that locally retains experimental phases while matching the predicted amplitude distribution. The statistical difference of voxel intensities between the predicted ensemble distribution and the baseline reference then allows computing a probability-derived confidence measure in the form of a rescaled z-score that we term the predicted voxel-wise difference test (pVDDT) score. Since Fourier amplitudes have been spectrally matched in baseline and feature-enhanced maps, the intensity residuals are dominated by phase errors in the predicted map (see Methods) and the pVDDT metric captures these errors in an interpretable way (Figure 5a, Supplementary Figure S6l-k).

**Fig. 5.**
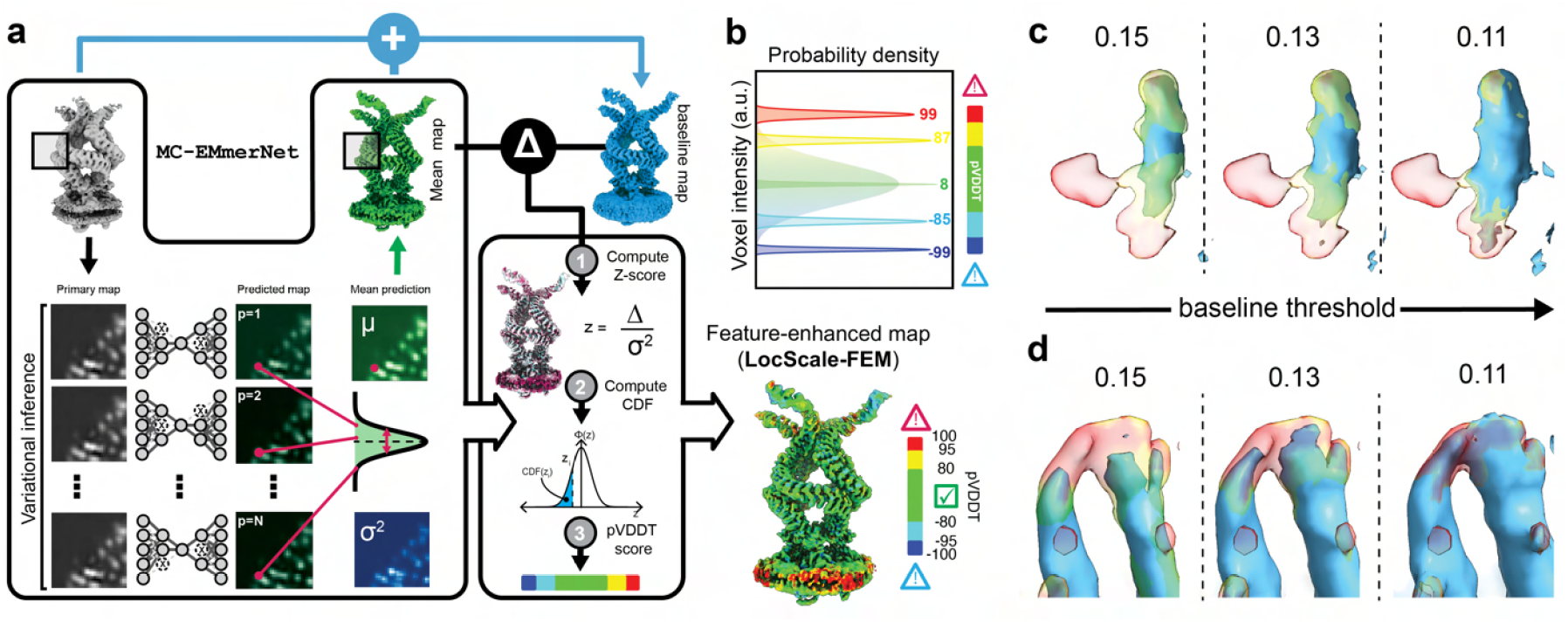
Confidence-weighted map optimisation and feature-enhanced maps (LocScale-FEM) (a) Schematic overview of Bayesian-approximate variational inference in MC-EMmerNet (left), and computation of voxel-wise pVDDT confidence scores by statistical comparison to a baseline reference (right). The far right shows the resulting feature-enhanced map (FEM) weighted according to local confidence. (b) Illustration of predicted and scaled central voxel intensity distributions for five voxels with varying pVDDT scores. For clarity, only a single predicted distribution is shown; in practice, unique distributions are generated per voxel. (c-d) Comparison of baseline and LocScale-FEM maps for plant phytochrome A (EMDB: EMD-35691). Shown are two regions with positive pVDDT scores (threshold: 0.15), overlaid with baseline maps displayed at varying thresholds, with (c) showing an example where the baseline map nevers recovers the positively flagged density. See also **Supplementary Figures** S6–S7.

### Interpreting pVDDT confidence scores

While the variances from Monte Carlo dropout provide a first estimate of predictive uncertainty, we found that they were systematically smaller than the actual prediction errors, measured as the absolute difference between the predicted means and the corresponding baseline values (Supplementary Figure S6h). This mismatch reflects the common tendency of Monte Carlo dropout to underestimate uncertainty and produce overconfident predictions ^42,43^. We therefore used isotonic regression calibration to adjust the predicted standard deviations so that they matched the empirically observed residuals (Supplementary Figure S6i). The resulting calibrated uncertainties establish a statistically robust baseline for voxel-wise pVDDT scores, which quantify both the probability and direction of signal deviations in the predicted maps (Figure 5b-c, Supplementary Figure S6l-m). The magnitude of the score reflects the statistical strength of the deviation. For interpretability, we define three ranges of pVDDT values. Scores between –80 and +80 fall within the reliable range, corresponding approximately to the central 80% confidence interval of the standard normal distribution. This threshold provides a practical compromise between sensitivity and robustness of the metric. Scores below –80 or above +80 indicate statistically significant deviations, while values beyond ± 95 mark voxels where deviations are highly unlikely given the calibrated uncertainty. As feature-enhanced maps tend to accentuate secondary structure, negative pVDDT confidence scores often appear along the grooves of *α*-helices. High positive pVDDT scores (greater than +95) may represent network hallucination of density features not present in the phase-conservative baseline map (Figure 5c, Supplementary S7a), but in other cases may correspond to real features (Figure 5d, Supplementary S7b). Thus, pVDDT confidence scores help draw attention to regions that merit closer inspection, and joint examination of feature-enhanced and baseline maps provides an efficient strategy for guiding interpretation. One limitation of our procedure is that pVDDT values are referenced to the feature-enhanced maps for visualisation; as a result, strongly negative scores corresponding to genuine densities that are completely suppressed in the prediction will not be reported. Feature-enhanced maps should therefore always be evaluated alongside the baseline map to ensure that such features are not overlooked.

### Applications of LocScale-FEM in structure analysis

We examined how pVDDT confidence scores can guide interpretation of feature-enhanced maps from LocScale-FEM. As a first example, we applied LocScale-FEM to the unmodified half maps of EMD-13234 to re-examine the *in-cell* subtomogram averaged reconstruction of M. pneumoniae 70S ribosomes ^8^. For comparison, we also generated the corresponding map using DeepEMhancer, which has been trained on LocScale-1.0 (model-based) maps ^30^. The LocScale-FEM map effectively recovered weak density at levels comparable to the equivalent hybrid LocScale-2.0 map (cf. Supplementary Figure S4a-c), including density at the mRNA entry site, A-site tRNA and the nascent peptide (Figure 6a). pVDDT confidence scores supported the reliability of most of these features, with the exception of the nascent peptide density ^8^. While density near the mRNA entry site remained heterogeneous and uninterpretable, the A-site density in the LocScale-FEM map supported confident tRNA placement (Figure 6b). In contrast, DeepEMhancer maps showed pronounced downscaling of the same regions, leaving them essentially uninterpretable (Figure 6a,b).

**Fig. 6.**
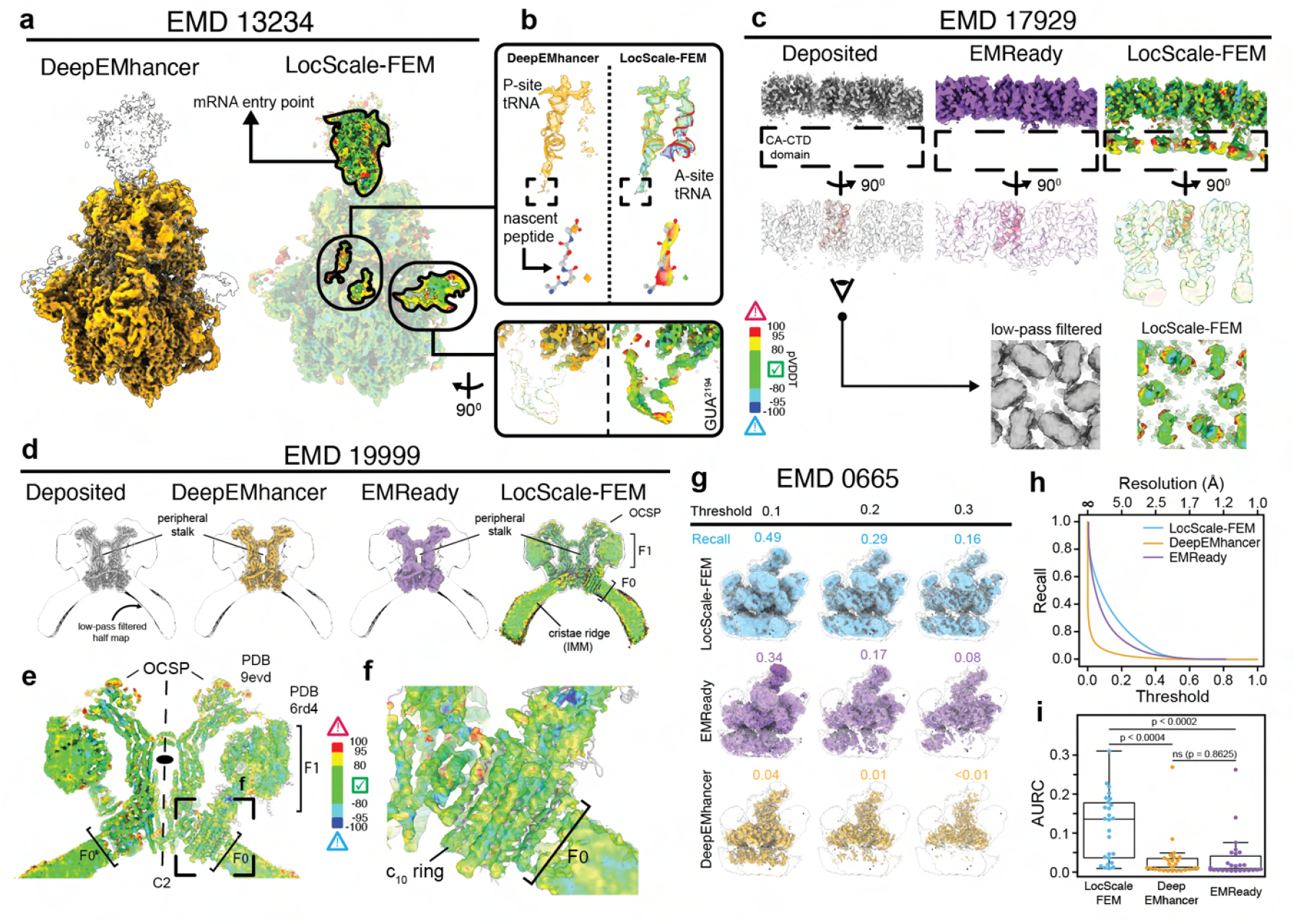
Comparison of LocScale-FEM and alternative deep learning approaches for contextual structure recovery. (a) LocScale-FEM and DeepEMhancer maps of *Mycoplasma pneumoniae* 70S ribosome (EMDB: EMD-13234). Selected contextual densities are highlighted in the LocScale-FEM map. The 1% FDR contour computed from the unmodified half maps is outlined for both maps. (b) Close-up views from (a), showing improved interpretability of densities for the A-site tRNA, nascent peptide, and a peripheral region. (c) Comparison of author-deposited, EMReady-optimised, and LocScale-FEM maps for subtomogram averages of the Human T-cell leukemia virus type I immature capsid (CA;EMDB: EMD-17929). The capsid protein C-terminal domain (CA-CTD) is highlighted. A close-up shows the CA-CTD viewed from below, comparing the 25 Å low-pass filtered EMDB map with the LocScale-FEM map. (d) Author-deposited and optimised maps for in situ subtomogram averages of the peripheral stalks of *Polytomella* ATP synthase (EMDB: EMD-19999). All optimised maps are superimposed on the low-pass filtered contour of the EMDB-deposited map. Contextual densities for the F_1_ and F_0_ ATP synthase subunits, oligomycin sensitivity-conferral protein (OCSP), and the crista membrane are indicated. (e) Close-up of the *C*2-symmetric ATP synthase dimer in the LocScale-FEM map. The right half of the map is shown transparent and superposed on PDB 6rd4 (F_1_ and F_0_) and PDB 9evd (peripheral stalk). The pVDDT colour bar is shown for reference. (f) Zoom of the F_0_ subunit showing resolved helical densities for the *C*_10_ ring. (g) Volume recall for EMD-0665 computed for different thresholds for LocScale-FEM, EMReady and DeepEMhancer optimised maps. (h) Volume recall curves for LocScale-FEM, DeepEMhancer and EMReady maps generated for EMD-0665 (i) Area-under-the-recall-curve (AURC) for LocScale-FEM, DeepEMhancer and EMReady plotted for 26 randomly selected maps from the EMDB. Statistical significance of mean differences was evaluated by permutation testing with 10,000 resamples. See also **Supplementary Figure** S7.

A similar trend was observed for subtomogram averages of tubular lattices formed by the capsid (CA) and nucleo-capsid (NC) domains of Human T-cell leukemia virus type I (EMD-17929) ^48^. LocScale-FEM produced interpretable density for dimers within the the irregular hexamers formed by the CA C-terminal domain (CA-CTD), which in the EMDB-deposited map is visible only after strong low-pass filtering (Figure 6c; shown at 25 Å low-pass) and entirely suppressed in the corresponding EMReady map. It has been hypothesised that CA-CTD interactions buttress the NTD scaffold but leave the lattice free to adopt diverse curvatures ^48^, highlighting that co-visualisation of ordered elements (here, the NTD lattice) and less-ordered components (here, CTD-mediated contacts) in such reconstructions may help inform mechanistic models.

Another class of structures for which co-visualisation of contextual density is highly informative are transmembrane or membrane-associated complexes. An illustrative case is the *in-cell* structure of mitochondrial ATP synthase from *Polytomella* ^49^. The deposited consensus map (EMD-19999) captures ATP synthase dimers on cristae ridges, with the best resolved region (4 Å local resolution) at the central peripheral stalk. The F1 and Fo subcomplexes are less well resolved due to averaging over different rotational states ^49^, but remain visible after low-pass filtering (Figure 6d). Map optimisation with DeepEMhancer and EMReady entirely suppressed density outside the central peripheral stalk, whereas the LocScale-FEM map co-visualised the peripheral stalks alongside both F1 and Fo subcomplexes and the strongly curved cristae membrane, without loss of detail in the peripheral stalks compared to the deposited or otherwise density-modified maps. The LocScale-FEM map also distinguished individual *α*-helical densities for several of the ten outer c-ring subunits embedded in the membrane (Figure 6e,f), with pVDDT scores suggesting high confidence for these features. This case exemplifies how LocScale-FEM can preserve the broader architectural context of complex membrane-bound assemblies. Similarly, LocScale-FEM maps of the yeast COPII inner coat assembled on membrane tubules (EMD-15949) ^5,50^ allowed confident placement of the Sar1 amphipathic helix inserting into the outer leaflet of the membrane (Supplementary Figure S7d). The map clearly visualises the parallel insertion of the amphipathic helices, which has been posited to reinforcing curvature through cooperative interactions across the lattice ^51^. Additional examples are provided in Supplementary Figure S7e-f.

The above examples suggest that LocScale-FEM is less prone to inadvertent signal suppression, preserving contextual density features that are often obliterated by other deep-learning–based map optimisation methods. To quantify this effect, we measured volumetric recall relative to an unbiased signal envelope derived from the original half maps using FDR thresholding ^52^. For each case, LocScale-FEM, DeepEMhancer, and EM-Ready maps were normalised by rescaling their positive voxels to the unit interval [0,1]. We then calculated volumetric recall across varying thresholds to generate recall curves for each method. Figure 6d,e illustrates an example for this procedure. Applied to 26 randomly selected EMDB reconstructions, LocScale-FEM showed significantly higher integrated volume recall (AURC) compared to DeepEMhancer and EMReady, confirmed by permutation testing (p < 0.0004 and p < 0.0002, respectively; Figure 6f). This analysis establishes that the advantages observed in individual examples generalise across diverse datasets.

### Interpretation of small-molecule ligands

We next examined the utility of feature-enhanced maps and pVDDT scores for interpreting small-molecule ligands and other non-protein components. As a representative case, we analysed the GPCR–Gi complex of human relaxin family peptide receptor 4 (RXFP4) bound to compound 4, an amidrazone scaffold (EMD-33888) ^53,54^. LocScale-FEM reproduced the ligand density observed in the deposited (DeepEMhancer-processed) map, with visible features corresponding to both the hydroxyindole and chlorophenyl substituents (Figure 7a). Analysis of pVDDT confidence scores, however, revealed important distinctions between these moieties. While the scores supported reliable interpretation of the hydroxyindole group, strongly positive scores around the chlorophenyl substituent indicated possible overemphasis of density at this site relative to an amplitude-modified baseline. This differential confidence scoring demonstrates how our approach can provide additional guidance for ligand interpretation by flagging regions where structural features may be less reliable or potentially artifactual. To validate this approach across different ligand chemistries, we examined β-galactosidase bound to the inhibitor phenylethyl-β-D-thiogalactopyranoside (PETG; EMD-7770) ^55^, where ligand pose can be challenging ^56^. The LocScale-FEM analysis showed high confidence density for the well-ordered pyranose moiety while appropriately downscaling density for the somewhat more flexible phenylethyl substituent (Supplementary Figure S8b). This pattern suggests that our confidence scoring effectively distinguishes between well-ordered structural elements and regions with inherently higher mobility or weaker density. LocScale-FEM demonstrated density preservation across diverse ligand types and retained ligand density also in cases where other deep-learning methods tend to downscale density excessively (Figure 7b,c, Supplementary Figure S8c-e). Despite this apparent robustness, fundamental limitations remain. Ligands are strongly underrepresented in the EMDB and their structural diversity spans an enormous chemical space, which places inevitable constraints on any deep-learning model’s ability to generalise across all molecular scaffolds. This limitation is exemplified in our analysis of ordered lipid densities in the Connexin-43 gap junction channel (EMD-33394) ^57^ (Supplementary Figure S8a). While such densities are clearly visible in the baseline LocScale maps, they were substantially downweighted in LocScale-FEM as well as in DeepEMhancer and EM-Ready predictions. This case shows that all three deep learning models may systematically suppress genuine signal. Taken together, these examples illustrate both the potential and the limitations of LocScale-FEM for non-protein components. Confidence scores can provide useful discrimination in some cases, while in others suppression of genuine features persists. In practice, joint inspection of feature-enhanced maps, confidence scores, and baseline references will therefore be essential for interpretation.

**Fig. 7.**
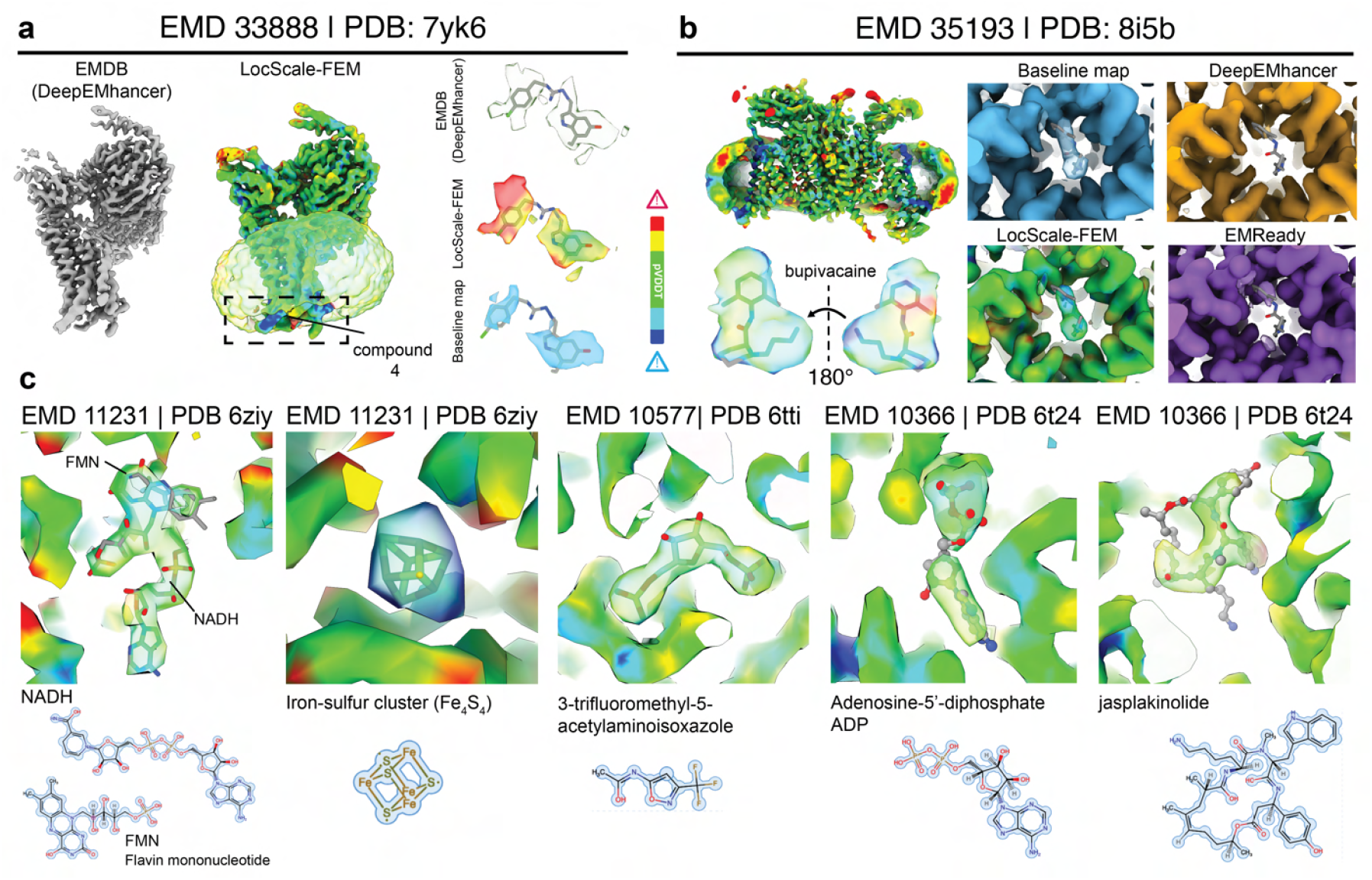
Small-molecule ligand interpretation with LocScale-FEM. (a) Author-deposited (DeepEMhancer) and LocScale-FEM map of (RXFP4)-G_i_ bound to 1-[2-(4-chlorophenyl)ethyl]-3-[(7-ethyl-5-oxidanyl-1H-indol-3-yl)methylideneamino]guanidine (compound 4; EMDB: EMD-33888). Ligands superposed on their respective densities are shown on the right. (b) LocScale-FEM map for human voltage-gated sodium channel (Nav1.7; EMDB: [EMD-351993](https://www.ebi.ac.uk/emdb/EMD-35193)) in complex with bupivacaine ^44^ The close-ups show ligand-superposed densities for LocScale-FEM, DeepEMhancer and EMReady-optimised maps. The LocScale-FEM baseline map is also shown for comparison. (c) LocScale-FEM maps for *Thermus thermophilus* complex I (EMDB: EMD-11231) ^45^, pyruvate kinase PKM (EMDB: EMD-10577) ^46^ and stabilised F-actin filaments (EMDB: EMD-10366) ^47^ containing a chemically diverse array of ligands. 2D chemical representations of the respective ligands are drawn below. See also **Supplementary Figure** S8.

## Discussion

In this work, we demonstrate new capabilities of LocScale-2.0 map optimisation for recovering weak and contextual structure in reconstructions affected by conformational or compositional heterogeneity, both from cryo-EM single-particle analysis and cryo-ET subtomogram averaging. LocScale-2.0 integrates physics-informed map sharpening with uncertainty-aware deep learning to address limitations of current map post-processing methods. Both hybrid sharpening and feature-enhanced maps generated by LocScale-FEM consistently mitigate negative bias in regions that remain poorly resolved in consensus maps or retain residual heterogeneity after extensive classification. These improvements are most pronounced for low-resolution regions and contextual density, while performance in the highest-resolution areas varies due to limitations of current physical priors and residual trade-offs between preserving low-resolution context and maximally accentuating high-resolution detail. Importantly, LocScale-FEM introduces calibrated voxel-wise confidence scores that enable systematic identification of ambiguous regions, thereby reducing reliance on experience and subjective judgment. Although not all retained low-resolution density will be chemically interpretable, the ability of LocScale-2.0 maps to co-visualise high-resolution details and weaker contextual features at a single density threshold represents an important practical advantage for structural analysis and the visual communication of complex macromolecular assemblies.

Several limitations warrant consideration. pVDDT confidence estimates are referenced to the feature-enhanced maps, and strongly negative deviations corresponding to genuine features completely suppressed during prediction may go undetected. This is most relevant for small-molecule ligands and other components under-represented in the training set, underscoring the need to inspect feature-enhanced maps alongside their baseline references. Likewise, while our physical priors capture general scattering behaviour, they do not account for all aspects of macromolecular structure and further improvements can be made. Finally, although throughout we refer to significant differences between predicted and baseline maps as phase errors in the context of confidence estimates, it should be emphasised that the experimental Fourier phases themselves are subject to error. Since the true, error-free structure is unknown, the metric should therefore be seen as a conservative measure erring on the side of caution. Put differently, it should be recognised that some apparent errors may reflect genuine improvements over noisy observations but cannot be distinguished as such.

Future extensions could refine physical priors through explicit anisotropy corrections to account for direction-dependent blurring, and incorporate expectations on map behaviour in real-space such as solvent flattening ^58–60^, histogram matching ^61,62^, and the use of symmetry-related correlations. Several of these ideas have been investigated in previous methods, but their combination remains unexplored ^26,27^. For LocScale-FEM, while Monte Carlo dropout offers a practical means to estimate uncertainty, Bayesian alternatives that are currently computationally too demanding may in the future yield better calibrated or more accurate uncertainty estimates ^63–65^.

By introducing a statistical measure of reliability, LocScale-2.0 represents a step toward automated cryo-EM map interpretation frameworks that transparently report on predictive uncertainty. More broadly, integrating confidence-aware optimisation into automated model building and refinement could increase accuracy and sequence recall of these methods toward lower resolution maps. As cryo-EM expands to increasingly complex systems, such approaches will be important for ensuring rigorous and objective structural analysis.

## ACKNOWLEDGEMENTS

We thank Keitaro Yamashita and Tom Burnley for valuable discussions, and Agnel Joseph for help with integrating LocScale-2.0 in CCP-EM Doppio. We are grateful to Ambroise Desfosses, Benedikt Junglas and the participants of the CCP-EM Icknield model building workshops for testing early versions of the method and for useful feedback. This work was supported by the European Research Council (ERC) under the European Union’s Horizon 2020 research and innovation programme (ERC-StG-852880 to AJ), the TU Delft AI initiative and the Collaborative Computational Project for Electron Cryo-Microscopy (CCP-EM). We also acknowledge the use of computational resources of the DelftBlue supercomputer provided by the Delft High Performance Computing Centre.

## DATA AND CODE AVAILABILITY

LocScale-2.0 is open-source software released under a BSD license and is available at https://github.com/cryotud/locscale, where detailed documentation is also provided. Pretrained MC-EMmerNet models (v0.3) can be accessed via Zenodo (DOI: https://zenodo.org/records/8211668). Scripts used to generate figures shown in this manuscript can be found at https://gitlab.tudelft.nl/aj-lab/publications/2025_LocScale-2.0.

## Methods

### Estimation of molecular envelope

The regions in a map that correspond to macromolecule and associated components (such as lipid, glycans or ligands) as opposed to solvent (i.e. the molecular envelope or boundary) are identified using False Discovery Rate-controlled statistical thresholding ^37^. Sampling windows located near the box edges are used to to estimate the background noise distribution; we note that the procedure will fail if the noise sampling box overlaps with part of the signal as can happen for filaments or subtomogram averages in which signal extends to (some of) the box edges. In these cases, noise center coordinates of sampling windows can be individually supplied. The size of the noise box is set to 20 pixels or 10% the unit cell dimension, whichever is higher. FDR confidence maps are binarised at 1% FDR to yield *confidence masks*.

### Estimation of total scattering mass

Using the molecular volume enclosed by the confidence mask, the number of required pseudoatoms is calculated assuming constant values for the mean protein density and the average atomic mass of the pseudoatoms:

The number of required pseudoatoms to fill this missing scattering mass is calculated assuming constant values for the mean protein density and the average atomic mass:

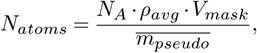

where *N*_*A*_ denotes Avogadro’s constant, *ρ*_*avg*_ (1.35 g/cm^3^) is the mean protein density determined from partial specific volumes using hydrodynamic compressibility ^66,67^ and *V*_*mask*_ is the volume of the FDR mask (for pure pseudo-models) or the difference mask (for hybrid models). The average atomic mass for pseudoatoms 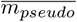 was estimated by computing the ratio of molecular mass to the number of non-hydrogen atoms over 119 atomic models from a reference dataset, which is derived from the training data used for DeepEMhancer. The median ratio was found to be 13.2 Da.

For hybrid model construction, we estimate the unmodelled volume by computing the difference between the FDR-based confidence mask and a binary mask derived from the available atomic model. The atomic model mask is obtained by placing a spherical mask of radius 3 Å at each atomic coordinate excluding hydrogen atoms. To minimise artefacts arising from subtraction, the difference mask is smoothed with a uniform filter across three voxels and subsequently binarised at a threshold of 0.5.

The calculated number of pseudoatoms are initially placed at random within the boundaries of the FDR confidence or difference mask. They are further randomly displaced by a magnitude of 0.5 pixels to prevent two atoms from overlapping.

### Positional refinement of pseudoatoms

The positions of pseudoatoms are then refined using a regularised gradient-based molecular dynamics scheme, with the cryo-EM density map acting as an external potential. This refinement integrates density-gradient forces and interatomic Lennard–Jones (LJ) interactions using a simple Verlet integration with friction damping ^68^. For simplicity, pseudoatom mass is set to unity, so that forces directly translate to accelerations. Note the atoms corresponding to the input atomic models remain unchanged during this operation.

The total force on a pseudoatom is computed as:

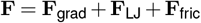

The attraction to high-density regions is modeled by the local gradient of the cryo-EM map:

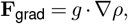

where *∇ρ* is the local gradient of the density map, and *g* is a scaling factor to scale the gradient force magnitude relative to that of the Lennard-Jones term (default *g* = 10). This approach is conceptually similar to other density-biased refinement methods ^69,70^.

To avoid pseudoatom clustering and promote spatial regularity, a pairwise Lennard–Jones potential is applied between pseudoatoms within a 3 Å neighborhood.To stabilise the refinement and assist convergence, we apply linear friction:

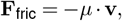

where **v** is the instantaneous velocity of the pseudoatom and *μ* = 10 is the friction coefficient. We implemented velocity Verlet integration to propagate the positions and velocities of pseudoatoms. By default. the refinement runs for 50 iterations with a timestep of *Δt* = 0.05 (in arbitrary units) which we found to reach convergence in most cases. Once the positions of all pseudoatoms are refined, pseudatoms are merged with the input atomic model to form a single structure. The pseudoatoms are introduced as a separate chain, with the sequence number of the added chain recorded for downstream purposes. For consistency, identifiers of the original atomic model chains always begin from “A”. If no input model is supplied, LocScale-2.0 assumes the entire map is unmodelled. Only the pseudoatomic model is retained for further processing. The overall workflow for constructing the pseudomodel and hybrid model is illustrated in Supplementary Figure S1a.

### ADP refinement of pseudoatoms and hybrid models

LocScale 2.0 performs ADP (isotropic *B*-factor) refinement with REFMAC (v5.8) ^71^ via the Servalcat wrapper (v0.4.100) ^72^, and supports explicit atomic models, pure pseudomodels, and hybrid models. In all cases positional refinement is disabled (`refi bonly`), all atomic *B* fac-^2^tors are initialised to 40 Å, and refinement is carried out against the unmodified map up to the Nyquist frequency.

REFMAC restrains ADPs using a weighted Kullback–Leibler divergence between atom pairs with the isotropic *B* factors as the optimised parameters, where the weights depend on distance and bonding connectivity ^71^. For hybrid models and pure pseudoatomic models, the absence of chemical bonds for the pseudomodel component necessitates a different strategy. Refinement is performed one cycle at a time, and each cycle is followed by spherical averaging of *B* factors within a 3 Å radius to enforce local smoothness and reduce the effective number of free parameters. In hybrid models, this averaging is restricted to the pseudoatomic chain(s) identified when constructing the hybrid model; coordinates belonging to explicit atomic chains are left unchanged except for their refined *B* factors. Pure pseudoatomic models lack valid chemical topology and are therefore refined in REFMAC with unconstrained refinement (`refi type unre`) while still applying the post-cycle spherical averaging step described above. This additional keyword is not required for hybrid models. By default, the sequence of a single refinement cycle followed by local spherical averaging is repeated ten times; the number of iterations can be adjusted by the user. Independent half map refinement is performed in all cases and automatically monitored for overfitting.

### Symmetry Averaging

If the input map has non-*C*1 point group symmetry, reference maps simulated from pseudo-models or hybdrid models do not necessarily preserve the point group symmetry of the original map. To correct for this, symmetry averaging on the simulated density using a user-defined point group symmetry. The desired point group symmetry can be specified using the `-sym` command-line argument.

### Amplitude scaling with generalised radial profiles

For each voxel position within the molecular envelope mask, cubic subvolumes are extracted both from the unmodified experimental map and from the corresponding reference map. The spherically averaged power spectra of these cubes are then compared, and the radial profile of the experimental cube is rescaled to match that of the reference cube. This procedure follows the principle of LocScale-1.0, but is extended to account for the use of pseudomodels and hybrid models.

For voxels that lie within the FDR or difference mask corresponding to pseudoatomic regions, additional corrections are introduced to incorporate average scattering profiles. First, the frequency limits for applying these corrections are defined: the low-resolution limit corresponds to the transition from a continuous-volume description to a discrete i.i.d. distribution, estimated from the local molecular volume ^35^, while the high-resolution limit is set by the half-map FSC. Within these limits, scattering amplitude deviations are computed by rescaling the scattering profile to the *B* factor of the reference profile, constructing an exponential fall-off function with the same *B* factor, and subtracting the scaled scattering amplitudes from this function. Finally, the computed deviations are added back to the reference radial profile within the relevant frequency range. This ensures that the local scaling not only matches the reference map but also reflects expected scattering behaviour in regions represented by pseudomodels.

### Monte Carlo (MC-EMmerNet)

#### Network architecture

We implemented a convolutional 3D U-Net, following a previous design ^30,73^ with several modifications (Supplementary Figure S6a; Table S6). The architecture consists of an encoder–decoder with skip connections, using strided convolutions for downsampling and transposed convolutions for upsampling. PReLU was chosen as the activation function throughout the network.

#### Dataset

The network was trained on cubic subvolumes extracted from average of unmodified EMDB-deposited half maps and their corresponding LocScale-2.0 reference maps. Cubes were 32 Å on each side and sampled from the training maps on a 24 Å grid. The dataset was derived from the DeepEMhancer map–model pairs ^30^, excluding cases where reliable confidence masks could not be generated. Hybrid model maps were available for 114 entries, of which 89 were used for training and 14 for validation (86:14 split). A separate set of 11 maps was reserved for testing and uncertainty calibration. In addition, 24 EMDB additional maps were specifically selected from the EMDB to assess the ability of LocScale-2.0 to enhance contextual density and retain non-protein components. These examples were also used to validate pVDDT scores (Supplementary Figure S7a–c,g–h).

#### Data augmentation

Training cubes were classified as signal-or noise-containing based on the resampled confidence mask. A 4:1 ratio of signal to noise cubes was enforced. Data augmentation included random rotations (0–2*π*), sharpening or blurring corresponding to random *B* factors between 0 and 400 Å^2^, and Gaussian low-pass filtering.

#### Training

Input intensities were standardised to zero mean and standard deviation of 0.1, and maps were resampled to 1 Å per pixel. The Adam optimiser was used with a constant learning rate of 10^−4^, and mean absolute error (MAE) served as the loss function. Networks were trained for 15 epochs.

#### Prediction

At inference, maps were preprocessed identically to training data. Subvolumes of size 32 Å were extracted every 16 Å and restricted to the confidence mask region. A default batch size of 8 per GPU was used, with multi-GPU data parallelism handled via `MirroredStrategy`.

Uncertainty estimates were obtained by enabling dropout during inference (`training=True`), yielding multiple stochastic forward passes. Voxel-wise means and standard deviations were computed across the ensemble. Overlapping cube predictions were merged using a weight matrix to reduce boundary artefacts. Final mean and standard deviation volumes were resampled to the original pixel size and saved as `.mrc` files.

### Uncertainty prediction

Multiple stochastic forward passes (MC samples) are performed to generate a distribution of predictions. From these, the voxel-wise mean and variance are computed as follows:

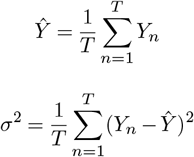

where *Ŷ* is the predictive mean, *σ*^2^ is the predictive variance, and *T* is the number of MC samples.

The mean and variance volumes are stored for each voxel in the map. During post-processing, these volumes are reintegrated into their original positions on the global grid and corrected for overlaps as described previously.

### Uncertainty calibration

The uncertainty estimates from Monte Carlo dropout are typically uncalibrated and therefore not directly suitable for statistical inference without further adjustment ^42,74^. In a perfectly calibrated regression model, the predicted uncertainties should match the expected residuals, defined as the absolute difference between the prediction and the true value. As shown in Supplementary Figure S6h, the raw uncertainties predicted via Monte Carlo dropout tend to be significantly smaller than the actual residuals.

For a given voxel *i*, the prediction interval is defined as:

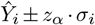

where *Ŷ*_*i*_ is the predicted mean, and *σ*_*i*_ is the standard deviation estimated via Monte Carlo dropout. The constant *z*_*α*_ is the z-score corresponding to a desired confidence level *α*.

A well-calibrated model produces uncertainty estimates that correspond to empirical probabilities. Specifically, for any confidence level *α* ∈ [0, 1], the probability that the true value *Y*_*i*_ lies within the interval defined by the predictive mean *Ŷ*_*i*_ and scaled uncertainty ± *z*_*α*_*σ*_*i*_ should be approximately *α*:

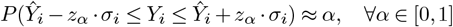

In this study, we employ isotonic regression to calibrate the predicted uncertainties ^43,75^ by learning a monotonically increasing function ℐ that maps predicted standard deviations to expected residuals.

The calibration procedure uses a set of *N* voxel-level predictions and their corresponding ground-truth values:

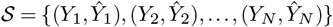

Residuals are computed for each voxel as:

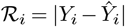

For the set 𝒰 = {*σ*_1_, *σ*_2_, …, *σ*_*N*_} of predicted standard deviations from Monte Carlo dropout, the isotonic regression function ℐ is learned by minimizing the squared error between residuals and calibrated uncertainties:

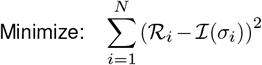

subject to: ℐ (*σ*_*i*_) ≤ ℐ (*σ*_*i*+1_) for all *i* ∈ {1, 2, …, *N* − 1}

The resulting function ℐ can then be applied to all predicted uncertainties to obtain calibrated standard deviations for statistical inference.

For the calibration procedure, 11 maps from the test dataset were selected. For each map, a mean and variance map was generated via Monte Carlo dropout. The mean map was used as input to generate the corresponding LocScale map, serving as the ground-truth reference. Calibration was performed voxel-wise by extracting the predicted mean and variance from the MC-EMmerNet predictions and the baseline values from the LocScale maps.

Since most voxels in a map correspond to background (i.e., regions with little or no signal), including them in the calibration process would bias the results and lead to underestimation of uncertainties. These background voxels typically exhibit very low predicted variance. However, simply discarding low-variance voxels is not trivial, as some signal-bearing voxels, particularly those with low intensity, also fall in this range. To address this, we apply a variance cutoff of 0.005: voxels with predicted variance below this threshold are excluded from the calibration process. While this introduces some miscalibration for low-intensity signal voxels, it significantly improves overall reliability. In practice, miscalibration tends to appear most prominently when low-threshold contours are used to visualize the feature-enhanced map. For consistent interpretation, we recommend using a contour threshold of 0.1 or higher when displaying predictions alongside pVDDT scores, as variances are well-calibrated in this range. Additionally, predicted variances are capped at a maximum value observed within the dataset. The isotonic regression model is also bounded accordingly: any input variance exceeding this cap is mapped to a fixed upper value. The effective calibration range is visualized in Supplementary Figure S6i.

Within this valid range, the calibrated variance serves as a reliable estimate of prediction error. This calibration is essential for the statistical interpretation of voxel-wise deviations and facilitates the detection of regions where the predicted distribution significantly diverges from the experimental reference. Such regions can then be analyzed using standard hypothesis testing techniques.

### Relationship of pVDDT score and phase differences between feature-enhanced and baseline maps

To evaluate voxel-wise significance of differences between predicted and baseline maps, we consider the data in Fourier space where density maps are represented by their series of Fourier coefficients. The structure factors of observed and predicted maps are:

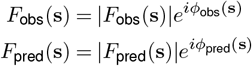

LocScale modifies the original observation by adjusting the amplitudes of its Fourier coefficients to match the radially averaged spectrum of the prediction while, within the scaling window, preserving their phases:

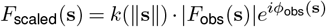

with:

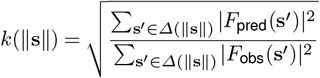

The local Fourier-space difference between the prediction and amplitude-scaled observation is:

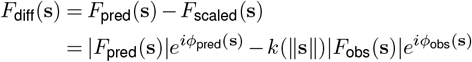

This can be rearranged to highlight the role of phase and amplitude differences:

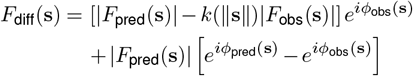

Here, the first term reflects residual amplitude differences between Fourier coefficients and the second term captures phase differences. When amplitude matching is effective (i.e., |*F*_pred_(**s**) | ≈ *k*(∥ **s** ∥) |*F*_obs_(**s**) |), the first term becomes small, and the difference density depends primarily on the phase difference (Supplementary Figure S6l,m):

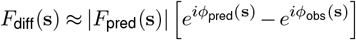

The inverse discrete Fourier transform over the local window grid 𝒢 gives the real-space difference density:

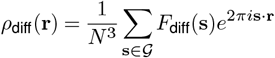

At the central voxel (**r** = 0):

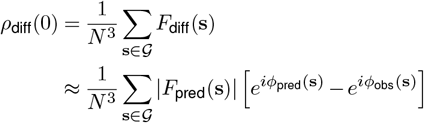

This demonstrates that, under the condition that amplitude differences are small, the central voxel intensity of the local difference map is primarily determined by the accumulated phase differences between Fourier terms of predicted and unmodified maps across spatial frequencies. These differences manifest as measurable voxel intensity variations in real space that can be statistically evaluated, and justify the use of the intensity difference of the central voxel *ρ*_diff_(0) as a phase-sensitive metric.

We use *ρ*_diff_(0), to compute a confidence metric based on a voxel-wise difference density test (*pVDDT*) score. A calibrated standard deviation *σ*_cal_ is first assigned to each voxel of the predicted map using a learned uncertainty from the Monte Carlo dropout predictions. The voxel-wise z-score for the difference between predicted and baseline map is then computed as:

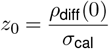

Here, *σ*_cal_ represents the expected difference between predicted and baseline densities. The z-score therefore measures how strongly the observed deviation exceeds this expectation.

The probability of observing a z-score less than or equal to *z*_0_ under the null hypothesis ℋ_0_ : *z*_0_ = 0 is given by:

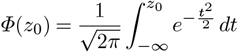

where *Φ*(*z*) is the cumulative distribution function of a standard normal distribution. To reflect both the magnitude and sign of the deviation in a bounded and interpretable scale, the result is linearly rescaled to the range [−100, 100]:

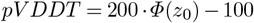

Positive pvDDT score indicate voxels where predicted voxel intensity exceeds the baseline reference (possible overestimation), negative values indicate underestimation, and larger absolute values reflect greater statistical significance.

### Automated model building with ModelAngelo

All ModelAngelo model building was done using ModelAngelo with default parameters. For each of the 50 map-model pairs in our ModelAngelo test dataset, FASTA files containing primary sequences of all complex components were pulled from the PDB. If applicable, protein and nucleic acid sequences were subsequently split over separate FASTA sequence files. For each model-map pair, ModelAngelo was run using both the EMDB-deposited map and LocScale-2.0 optimised map. To test the accuracy of sequence prediction, residue identities for each protein residue in the predicted structure were compared to that of the closest residue in the deposited PDB reference structure based on the coordinates of their respective C-*α* atoms. Recall statistics were then computed for all residues from each map-model pair in our dataset.

## Supplementary material

### Supplementary Figures

**Supplementary Fig. S1.**
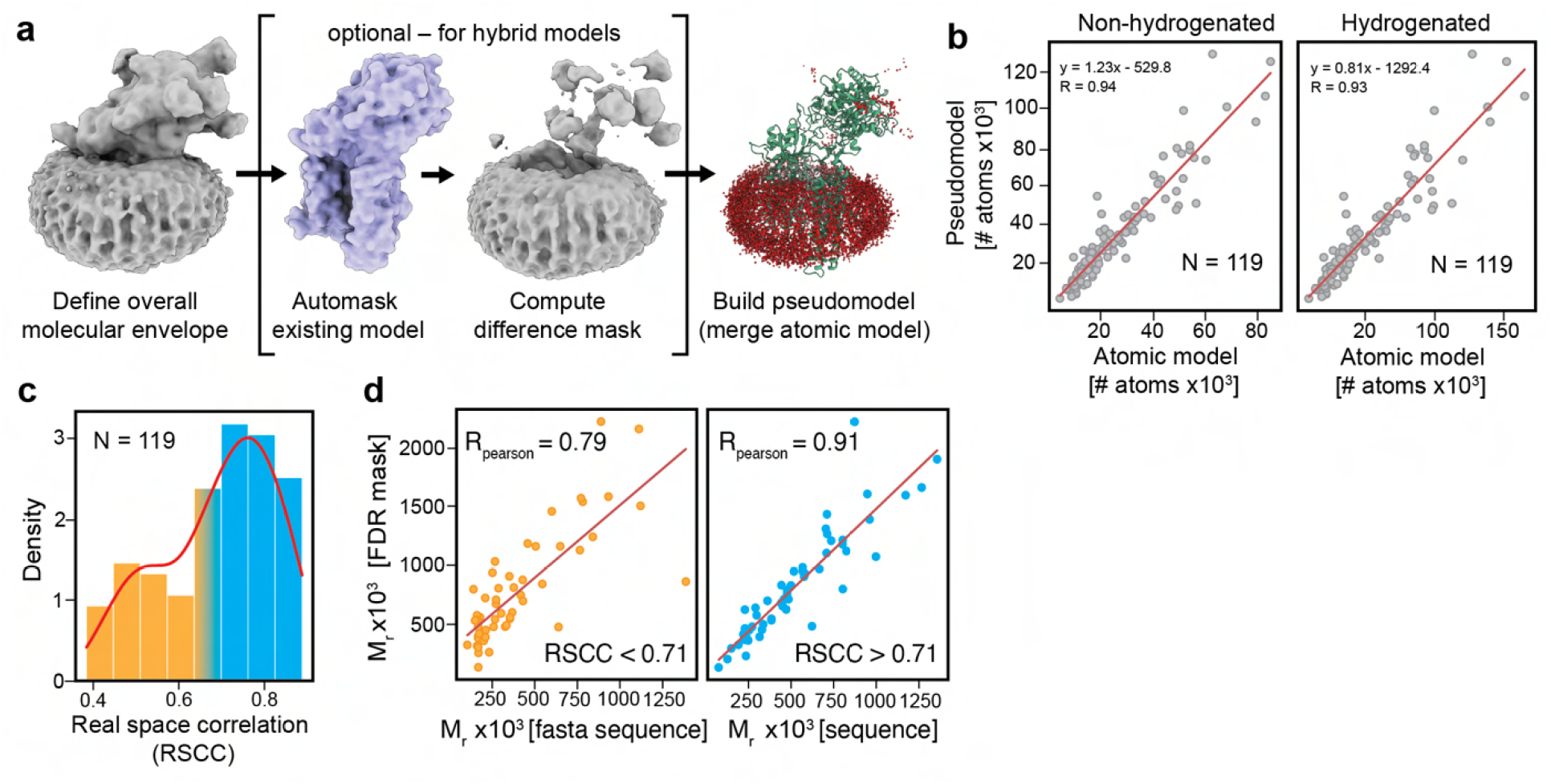
Definition of pseudoatom placement and generation of hybrid models. (a) Illustration of pseudoatom placement for human *γ*-secretase (EMD-3061). The molecular envelope is estimated using FDR thresholding ^52^, defined as the volume enclosed by a binarised mask computed from the confidence map at a 1% FDR threshold. When (partial) prior model information is available, a contextual structure mask is derived from the symmetric difference between the binarised model and envelope masks. Pseudoatoms are placed only within this difference mask and then merged with the prior atomic model. (b) Regression plots showing the number of pseudoatoms placed within the atomic model mask versus the expected number of atoms based on the known protein sequence, assuming hydrogenated and non-hydrogenated models. (c) Histogram of real-space correlation coefficients (RSCC) between FDR confidence masks and atomic model masks. Bars are colored by RSCC values below (RSCC < 0.71, orange) and above (RSCC > 0.71, blue) the median. (N = 119 for panels b–c.) (d) Regression plots comparing molecular mass estimates derived from pseudoatom placements within the FDR mask to those from the primary sequence, for RSCC below and above the median.

**Supplementary Fig. S2.**
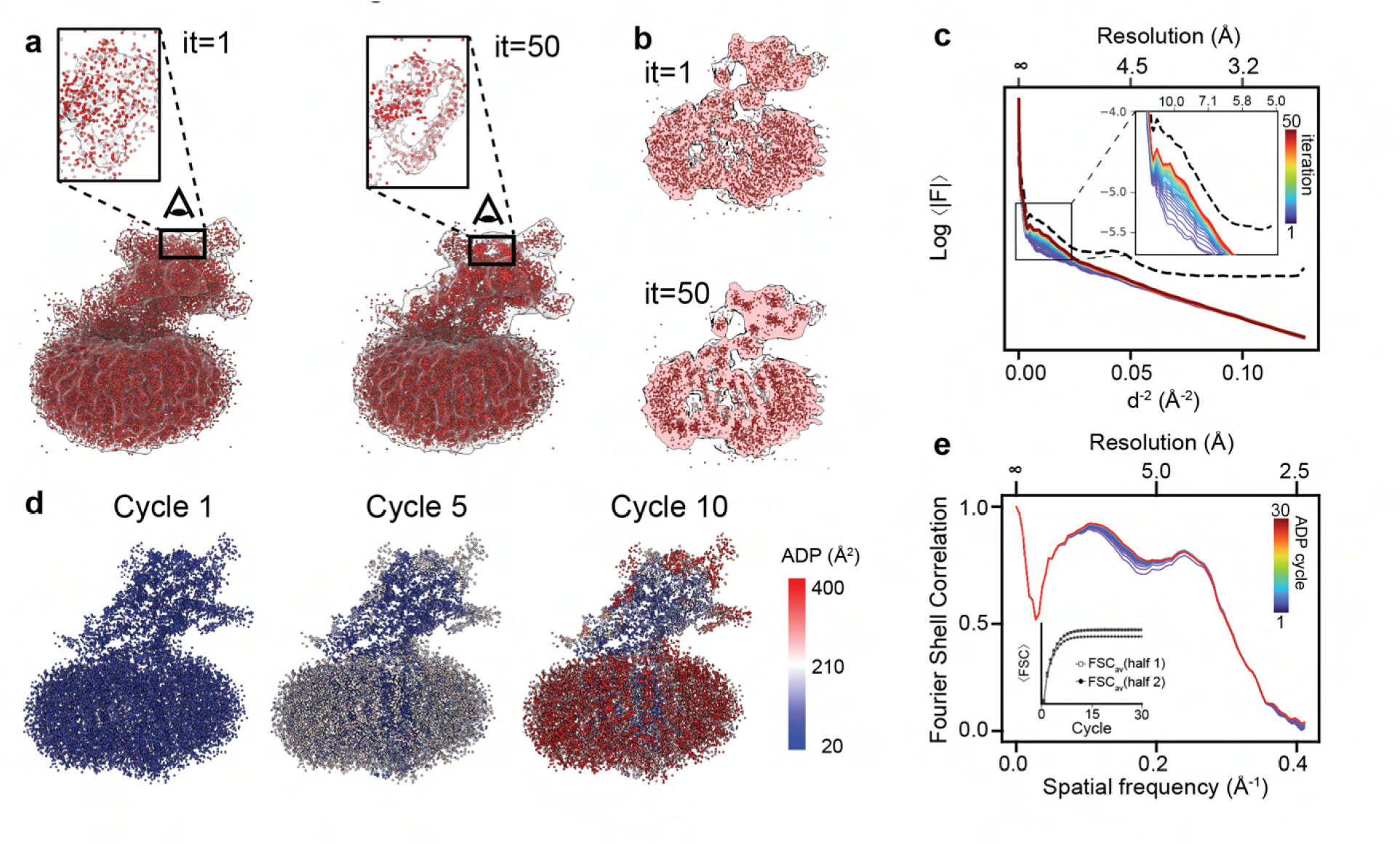
Pseudomodel generation and ADP refinement. (a) Example for pseudoatom placement and refinement for human *γ*-secretase (EMD-3061). Models are shown for the start (iteration 1) and end (iteration 50) of the positional refinement. The molecular boundary is outlined by a 1% FDR confidence mask. (b) Slab through the pseudomodel for both refinement iterations. (c) Radially averaged Fourier amplitudes of model maps computed throughout the iterations of pseudomodel refinement. Inset: Close-up of low-to-medium resolution region defining the overall molecular envelope. (d) Atomic displacement factors (ADP) mapped onto atomic coordinates shown for different snapshots across the ADP refinement cycles. (e) Evolution of model-to-map Fourier shell correlation (FSC) shown for a series of ADP refinement cycles. The insets shows the half-map validation displaying the average integrated FSC for work and test maps as a function of refinement cycle.

**Supplementary Fig. S3.**
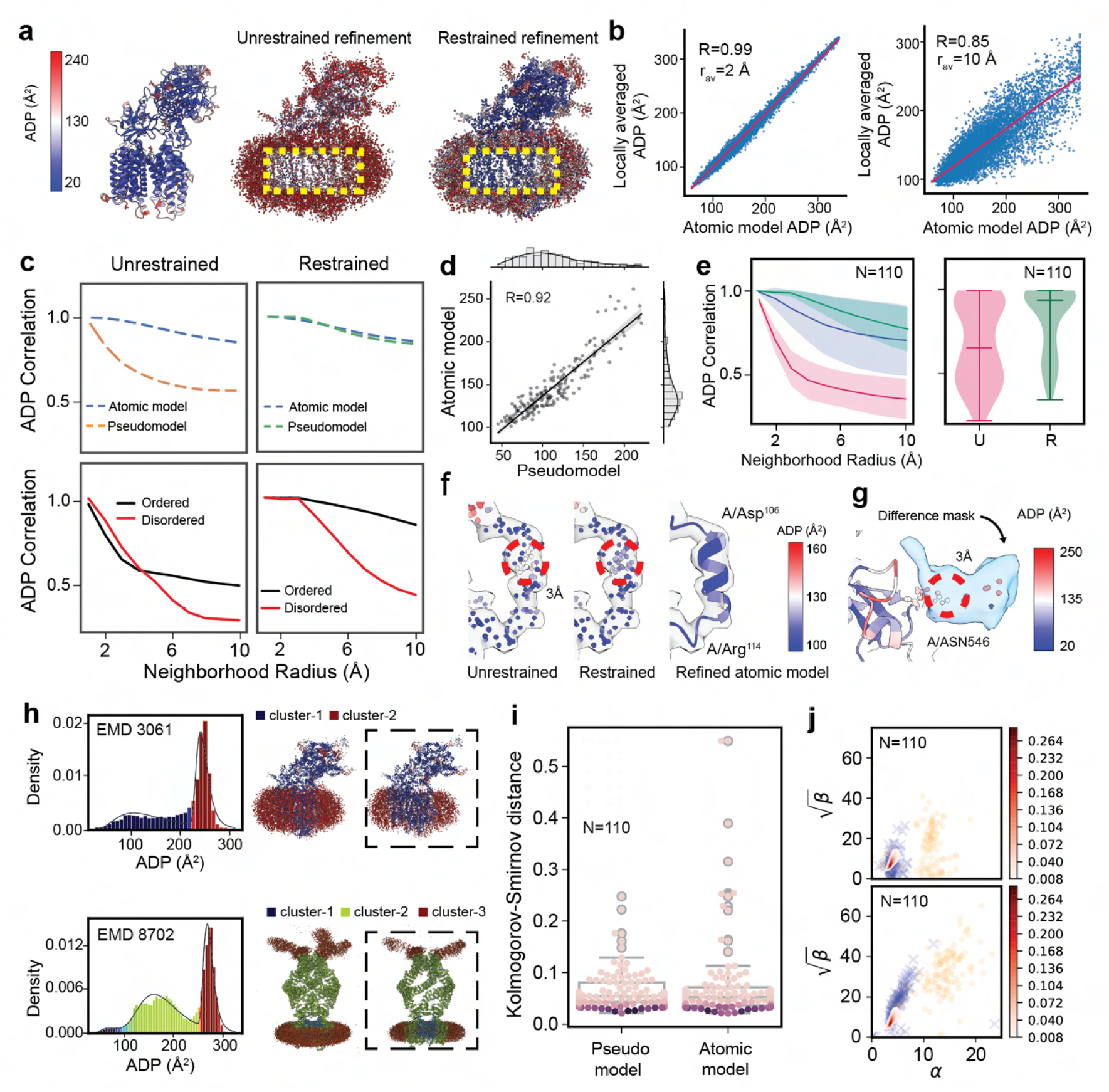
Implementation and validation of pseudomodel ADP refinement. (a) ADP distribution mapped on the refined atomic model of human gamma secretase complex (PDB ID 5a63). (b) Scatter plot of atomic ADPs versus locally averaged pseudomodel ADPs for averaging radii of 2 Å (left) and 10 Å (right). (c) Relationship between neighborhood radius and ADP correlation coefficient for unrestrained and restrained ADP refinement of pseudomodels compared to atomic model refinement (top). Also shown are neighborhood radius correlation curves for pseudoatom clusters in ordered and disordered regions of the map. (d) Regression plot of atomic model vs. pseudomodel ADPs for *γ*-secretase (EMD-3061/PDB ID 5a63). (e) Left: ADP correlation as a function of neighbourhood radius plotted for 110 randomly selected map/model pairs from the EMDB (red: unrestrained pseudomodel ADP refinement; green: restrained pseudomodel ADP refinement; blue: atomic model ADPs). Right: Similarity of the ADP correlation curves to a reference curve derived from an atomic model. (f) Effect of neighbourhood-restrained ADP refinement of pseudoatoms shown after four ADP refinement cycles. Red circles outline the 3 Å averaging radius (left and middle). For comparison refined ADPs of an explicit atomic model (PDB ID 5a63) are also shown for the same map segment (right). (g) ADP restraints at the boundary of explicitly modelled and pseudoatom regions in hybrid reference models. The difference mask outlines the unmodelled density of a N-linked glycan. (h) ADP validation with toBvalid ^39^ for pseudomodels obtained for EMD-3061 and EMD-8702. Histograms shows distinguishable modes of the shifted inverse *Γ* distributions (EMD-3061: two modes; EMD-8702: three modes). Mapping the different modes back onto the pseudomodel reveals dinstinguishable clusters corresponding to well-resolved and poorly resolved/disordered regions (right). (i) Plot of Kolmogorov-Smirnov distances measuring the maximum distance between the empirical cumulative distribution function of the pseudoatom or atomic model ADP distributions and the cumulative distribution function of the theoretical gamma distribution for 110 randomly selected maps/models from the EMDB. Statistical significance is obtained from permutation tests with 1000 resamples. (j) *α* versus *β*^1*/*2^ plots for bimodal fits to ADP distributions from atomic models (top) and pseudomodels (bottom). The red comet shows the smoothened unimodal distribution of *α* versus *β*^1*/*2^ from empirical analysis of 45,000 PDB structures ^39^.

**Supplementary Fig. S4.**
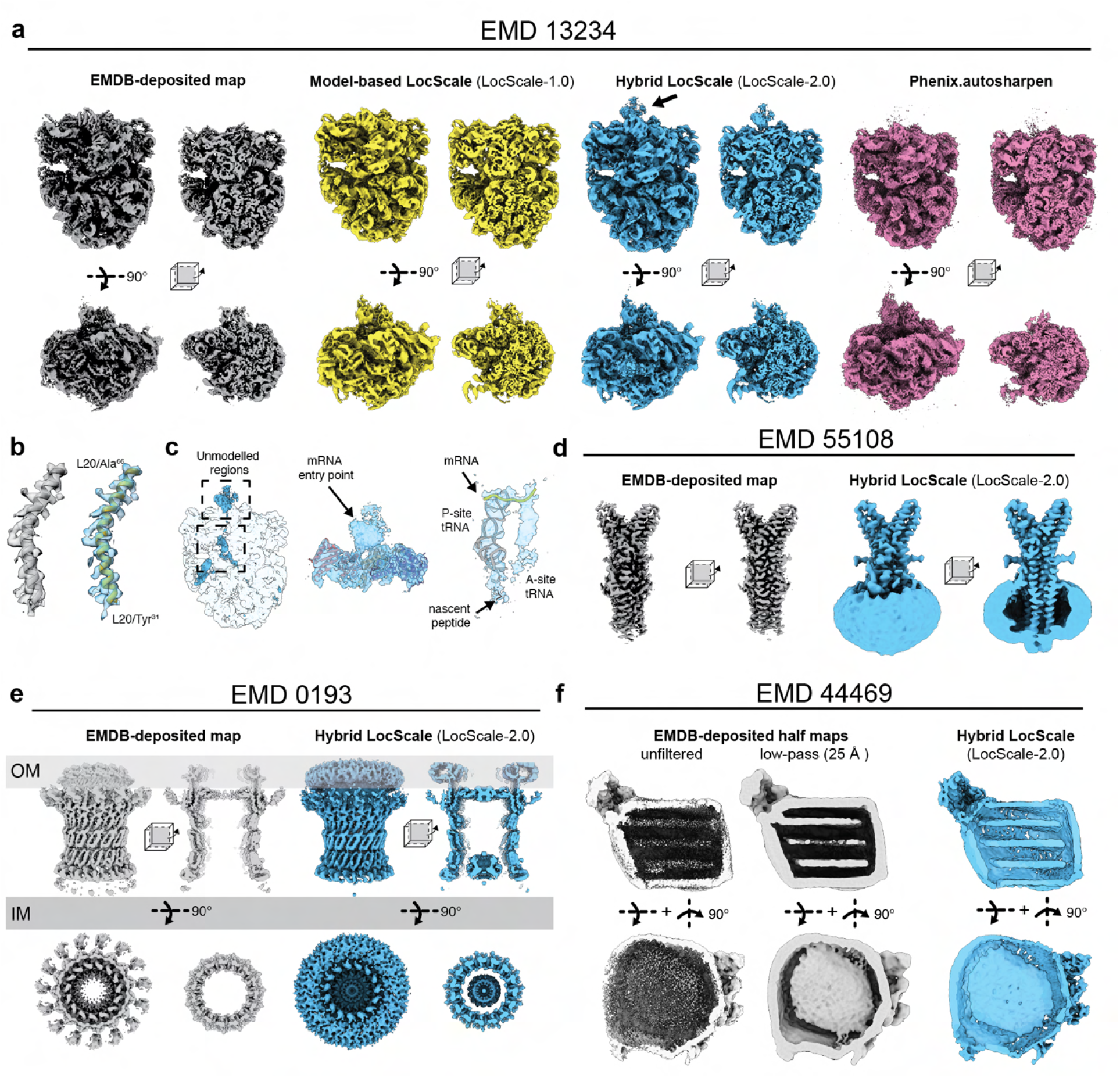
LocScale-2.0 map optimisation. (a) On-views and cross-sectional views of locally sharpened maps for in situ subtomogram averages of *Mycoplasma pneumoniae* 70S ribosomes (EMD-13234) using model-based sharpening with LocScale-1.0 (middle left), hybrid reference sharpening in LocScale-2.0 (middle right) and phenix.autosharpen (right). The author-deposited map is also shown (left). Arrows mark the mRNA entry point. (b) Side-by-side comparison of author-deposited and LocScale-2.0 maps for a well-resolved helical density segment from (a). (c) Unmodelled and partially interpretable regions of contextual structure highlighted in the LocScale-2.0 map. (d) Comparison of author-deposited and LocScale-2.0 sharpened maps for *Klebsiella pneumoniae* type II secretion system outer membrane complex (EMD-0193). (e) Comparison of author-deposited and LocScale-2.0 sharpened maps for Tweety homologue 2 (TTYH2, EMD-51108). (f) Unfiltered and low-pass filtered half maps in comparison to the hybrid LocScale map for ApoB100 (EMD-44469).

**Supplementary Fig. S5.**
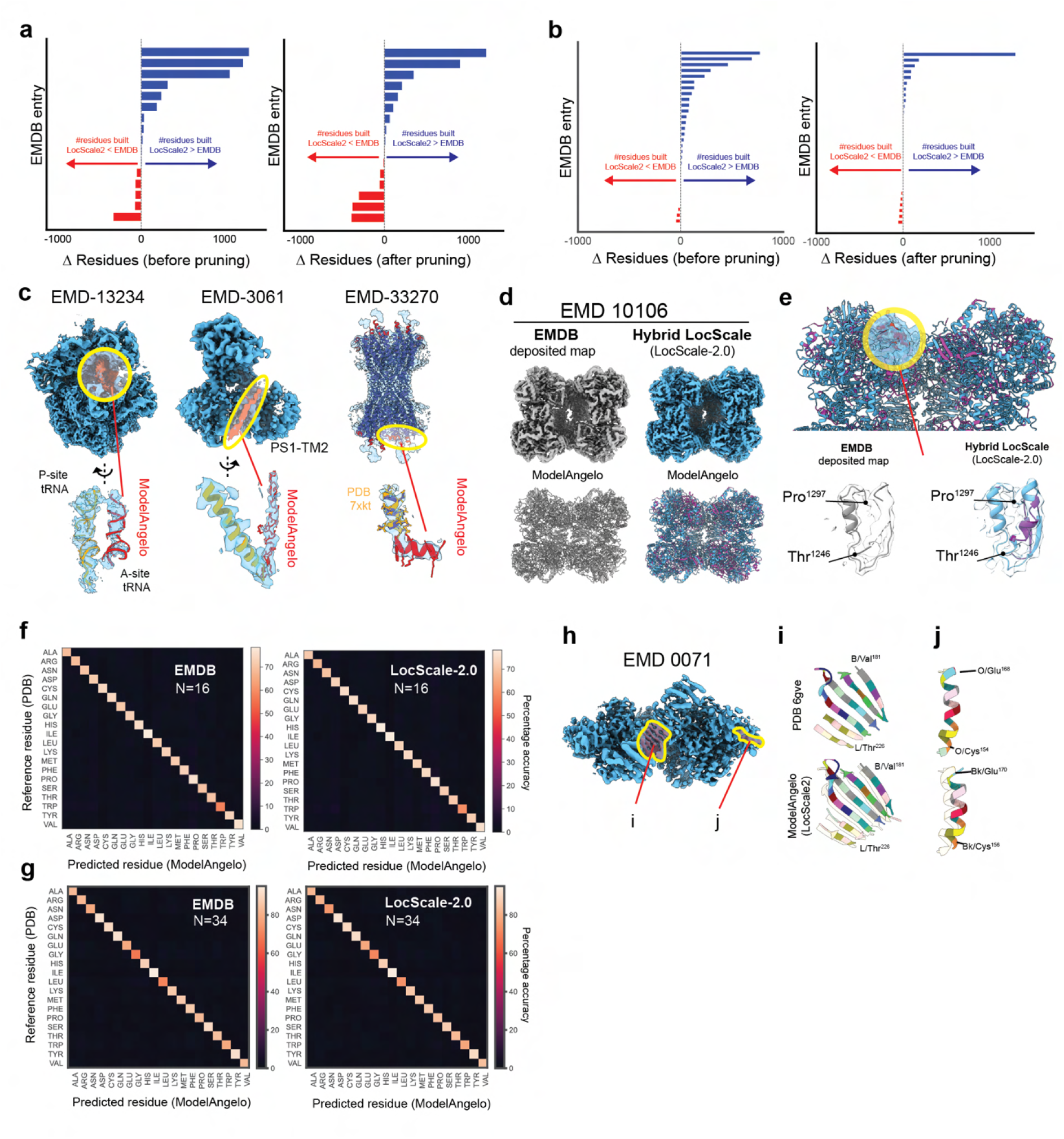
Automated model building with LocScale-2.0 maps. (a,b) Aggregate statistics of the number of ModelAngelo-built residues for 50 randomly selected maps from the EMDB before and after pruning; residue numbers are not normalised for point group symmetry. 16 of these models (a) were not part of the Modelangelo training set, and 34 had been used in ModelAngelo training (b). (c) Selection of EMDB entries for which LocScale-2.0 maps support improvements in ModelAngelo model completion compared to the deposited maps. Residues and nucleotide bases additionally built in LocScale-2.0 maps are shown in red. (d) LocScale-2.0 map for *Azospirillum brasilense* glutamate synthase (EMDB: EMD-10106). (e) Automatically built models from ModelAngelo superposed on a selected map region indicated in (a) using the author-deposited map (left) and LocScale-2.0 map (right) as input. The full map was used for automated model building. (f) Sequence recall for ModelAngelo predictions using deposited or LocScale-2.0 maps for 16 models not part of the ModelAngelo training dataset. (g) Sequence recall for ModelAngelo predictions using deposited or LocScale-2.0 maps for 34 models that had been used in the ModelAngelo training dataset. (h-j) Sequence recall visualised in EMDB: [EMD-0071/](https://www.ebi.ac.uk/emdb/EMD-0071)PDB: 6gve. Two representative regions are shown, one of which (j) shows a coordinate shift in the LocScale-2.0 map.

**Supplementary Fig. S6.**
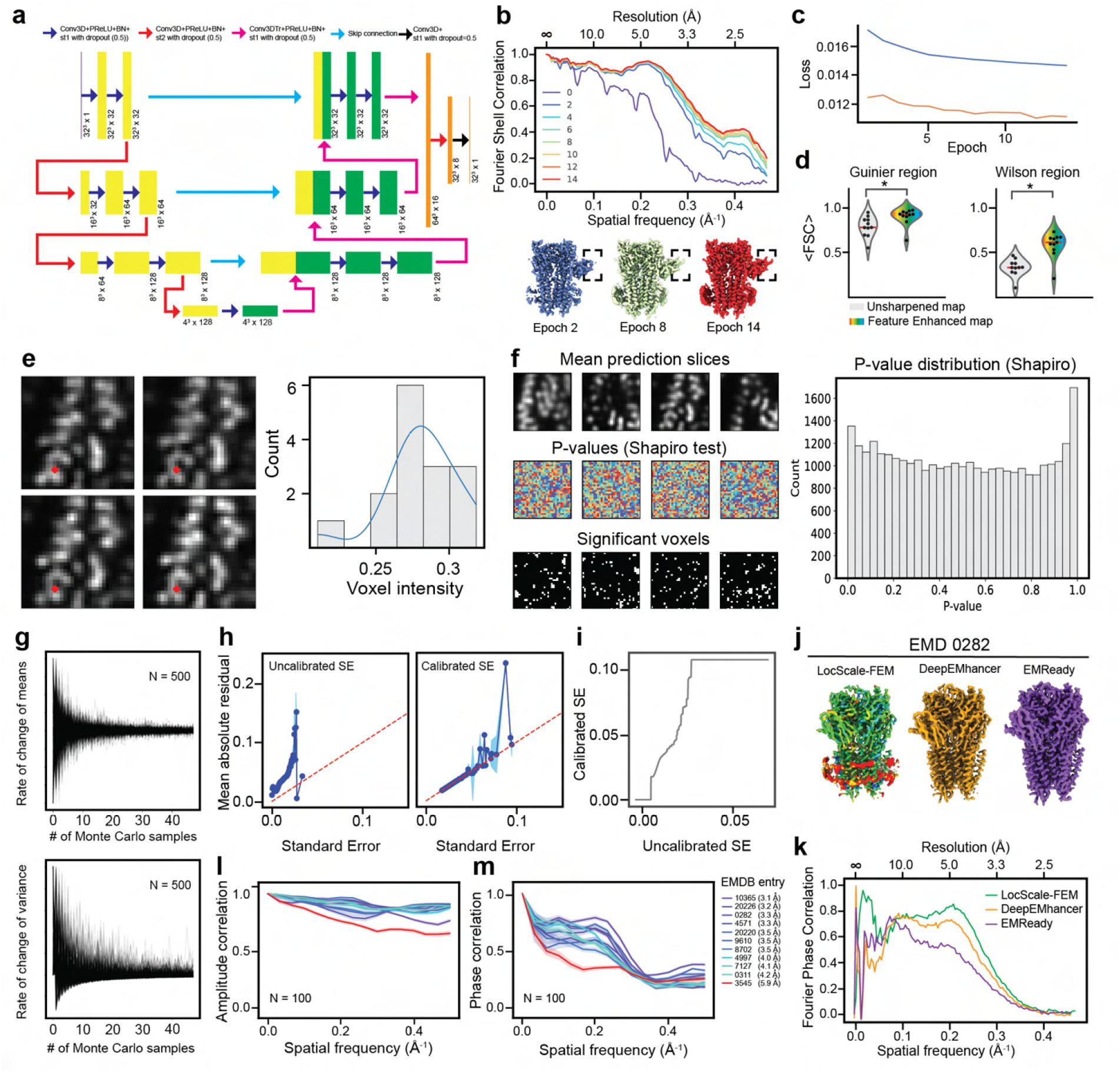
Monte Carlo Dropout and pVDDT scores. (a) 3D U-Net architecture of MC-EMmerNet. b) Spectral behaviour of MC-EMmerNet training measured by Fourier-Shell Correlation (FSC) of MC-EMmerNets test set prediction vs. target map as a function of training epoch. Low frequency information is rapidly learnt after a few epochs, while high frequency detail is learnt more gradually as observed by the slower convergence in the FSC curves at high frequencies. Predicted maps at selected epochs are shown. (c) Loss curves curves showing the evolution of training and validation losses. Due to the presence of dropouts in training the validation loss is lower than the training loss. (d) Average Fourier Shell correlation (<FSC>) in the Guinier region (0-0.1 Å^-1^) and Wilson region (>0.1 Å^-1^) measured vs. target map for the entire test set (N=11). (e) Central slices from 4 out of 15 samples generated by the MC-EMmerNet model using Monte Carlo dropout. The intensity distribution for the voxel highlighted in red is shown on the right. (f) Normality tests for all voxels in the sample cubes, with a histogram of the resulting p-values shown to the right. (g) Effect of the number of Monte Carlo samples on the change in prediction mean (top) and variance (bottom). (h) Reliability and calibration curves used to evaluate the uncertainty calibration. (i) Calibration curve from isotonic regression. (j) Feature-enhanced map (LocScale-FEM) for EMD-0282 colored by pVDDT scores, alongside DeepEMhancer and EMReady predictions. (k) Fourier phase correlation between the three deep learning models shown in (h) and the unmodified reference map. (l) Fourier amplitude correlation between predicted and target (hybrid LocScale-2.0) map for 11 EMDB entries with varying resolution between 3-6 Å. Shown are the mean correlations between 100 randomly selected signal cubes with their 95% confidence intervals. (m) Fourier phase correlation between predicted and unmodified (average of EMDB-deposited half maps) for 11 EMDB entries with varying resolution between 3-6 Å. Shown are the mean correlations between 100 randomly selected signal cubes with their 95% confidence intervals.

**Supplementary Fig. S7.**
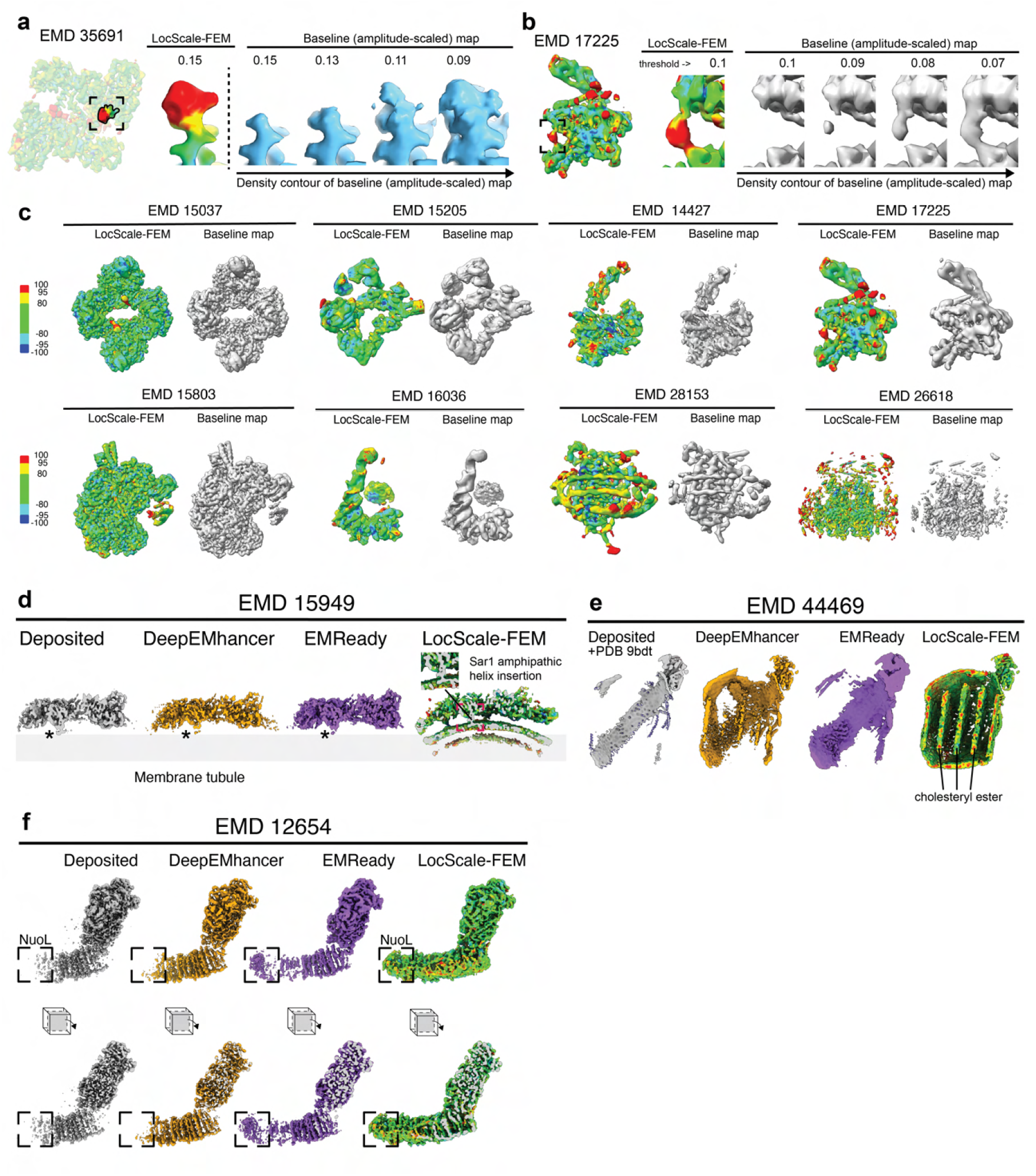
Interpretation of LocScale-FEM maps. (a) LocScale-FEM map for EMDB: EMD-35691) shown at 0.15 contour level. A feature with strong positive pVDDT score is shown as close-up. The corresponding baseline map of the highlighted feature displayed at successively decreasing contour levels is also shown. (b) LocScale-FEM map for EMDB: EMD-17225 shown at 0.15 contour level. A feature with strong positive pVDDT score is shown as close-up. The corresponding baseline map of the highlighted feature displayed at successively decreasing contour levels is also shown. (c) LocScale-FEM maps for eigth random EMDB entries shown alingsude their baseline maps. The maps are sorted according to increasing variance in the pVDDT scores. (d) Comparison of optimised maps for subtomogram averages of the yeast COPII inner coat assembled on membrane tubules (EMDB: EMD-15949). The inset shows density for the membrane-embedded amphipathic helix of the Sar1 subunit in the LocScale-FEM map. (e) Comparison of maps for Apolipoprotein B-100/low-density lipoprotein (LDL) particles EMDB: EMD-44469). Contextual densities for liquid-crystalline cholesteryl esters are highlighted. (f) On-axis and cross-sectional views of author-deposited and optimised maps for *E. coli* respiratory complex I (EMDB: EMD-12654. The peripheral antiporter subunit NuoL is highlighted.

**Supplementary Fig. S8.**
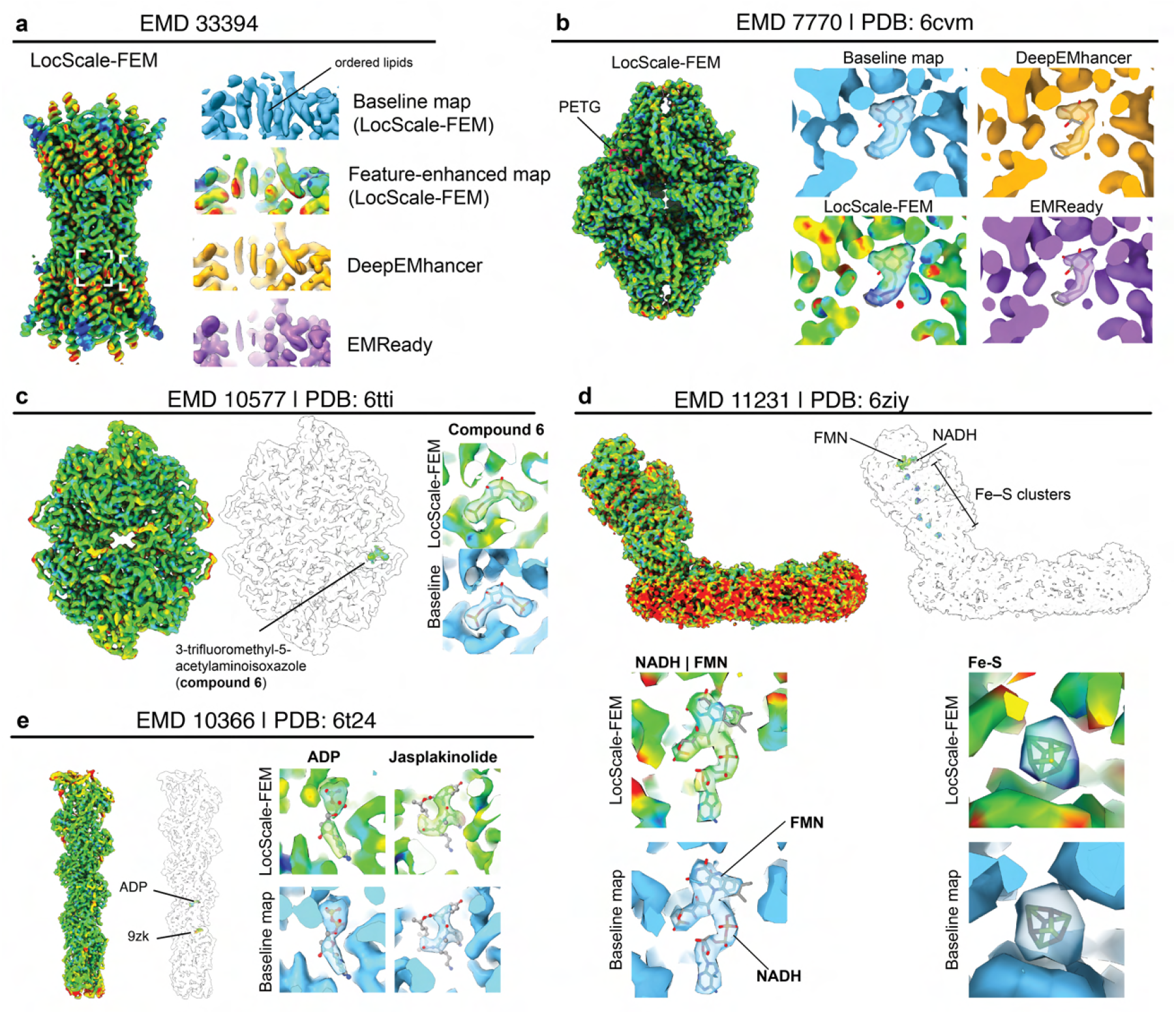
LocScale-FEM interpretation of small-molecule ligands. (a) Suppression of ordered lipid densities in Connexin 43 (EMDB: EMD-33394) in LocScale-FEM, DeepEMhancer and EMReady maps displayed at equivalent normalised thresholds. (b) Phenylethyl-β-D-thiogalactopyranoside (PETG) binding site in β-galactosidase (EMDB: EMD-7770) in LocScale-FEM, DeepEMhancer and EMReady maps displayed at equivalent normalised thresholds. PETG is shown in stick representation. (c) LocScale-FEM map of fragment-complexed human pyruvate kinase PKM (EMDB: EMD-10577). Close-ups show LocScale-FEM feature-enhanced and baseline maps of the ligand binding site. (d) LocScale-FEM map of fragment-complexed *T. thermophilus* complex I (EMDB: EMD-11231). Close-ups show LocScale-FEM feature-enhanced and baseline maps of the flavin mononucleotide/NADH site and an F_4_ S_4_ iron-sulfur cluster, respectively. (e) LocScale-FEM map of jasplakinolide-stabilised F-actin (EMDB: EMD-10366). Close-ups show LocScale-FEM feature-enhanced and baseline maps of the ADP site and an jasplakinolide binding sites, respectively.

### Supplementary Tables

**Supplementary Table S1.** Computation time for Hybrid LocScale-2.0 and LocScale-FEM optimisation.

|  | **EMD-3061** | **EMD-8702** | **EMD-44469** |
| --- | --- | --- | --- |
| Shape | 180×180×180 | 400×400×400 | 336×336×336 |
| 1% FDR volume [Å<sup>3</sup>] | 2.5 × 10<sup>5</sup> | 1.9 × 10<sup>6</sup> | 5.3 × 10<sup>6</sup> |
| **Hybrid LocScale** |  |  |  |
| Number of pseudoatoms | 9284 | 30887 | 206162 |
| Number of atoms | 10003 | 41900 | 38899 |
| Pseudo-model generation [min] | ~4 | ~18 | ~105 |
| ADP refinement [min] | ~14 | ~72 | ~117 |
| **LocScale-FEM** |  |  |  |
| Prediction [min]<sup>§</sup> | ~3 | ~10 | ~14 |
| **Summary** |  |  |  |
| Factor speed-up | 6 | 9 | 16 |

<sup>§</sup> LocScale-FEM predictions were done on a single Titan X (Pascal) GPU with 12 GB memory

**Supplementary Table S2.**
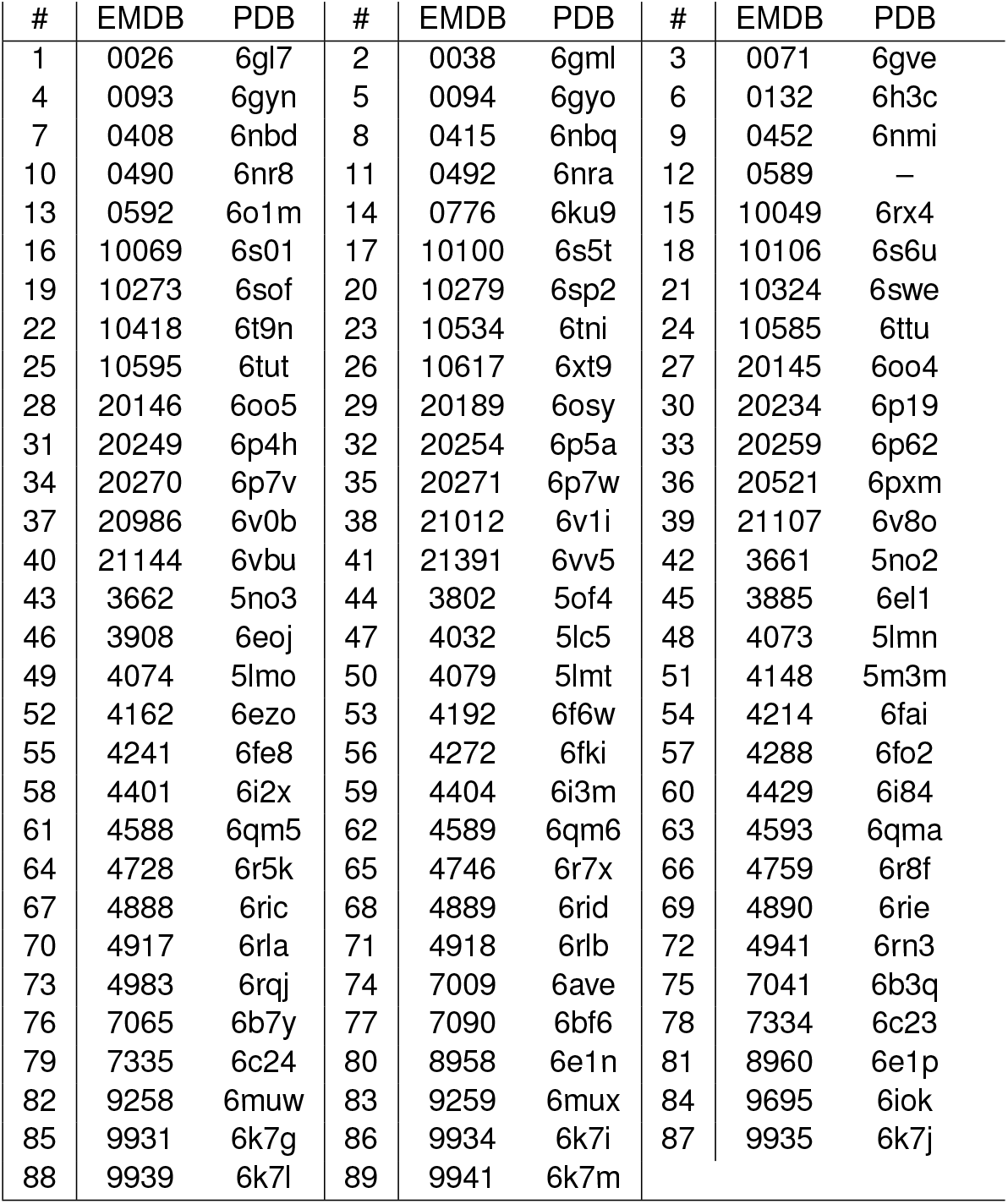
EMDB and PDB IDs for maps and models used for MC-EMmerNet training.

**Supplementary Table S3.**
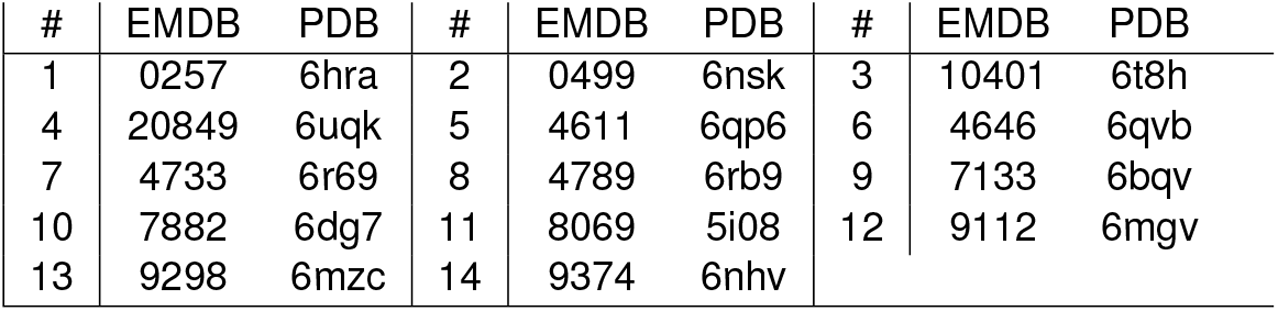
EMDB and PDB IDs for maps and models used for MC-EMmerNet validation.

**Supplementary Table S4.** EMDB and PDB IDs for maps and models used for testing and uncertainty calibration.

| # | EMDB | PDB | # | EMDB | PDB | # | EMDB | PDB |
| --- | --- | --- | --- | --- | --- | --- | --- | --- |
| 1 | 0282 | 6huo | 2 | 0311 | 6hz5 | 3 | 10365 | 6t23 |
| 4 | 20220 | 6oxl | 5 | 20226 | 6p07 | 6 | 3545 | 5mqf |
| 7 | 4571 | 6qk7 | 8 | 4997 | 6rtc | 9 | 7127 | 6bpq |
| 10 | 8702 | 5vkq | 11 | 9610 | 6adq |  |  |  |

**Supplementary Table S5.** EMDB and PDB IDs for maps and models in additional test dataset.

| # | EMDB | PDB | # | EMDB | PDB | # | EMDB | PDB |
| --- | --- | --- | --- | --- | --- | --- | --- | --- |
| 1 | 0193 | 6hcg | 2 | 51108 | 9g71 | 3 | 44469 | 9bdt |
| 4 | 35691 | 8isi | 5 | 13234 | 7p6z | 6 | 19999 | 9evd |
| 7 | 17929 | 8pu6 | 8 | 0665 | 6oa9 | 9 | 10106 | 6s6u |
| 10 | 33270 | 7xkt | 11 | 3061 | 5a63 | 12 | 17225 | 8ovx |
| 13 | 15037 | 7zz8 | 14 | 15205 | 8a64 | 15 | 14427 | 7z0l |
| 16 | 17225 | 8ovx | 17 | 15803 | 8b1r | 18 | 16036 | 8bgu |
| 19 | 28153 | 8ehw | 20 | 26618 | 7una | 21 | 15949 | — |
| 22 | 12654 | 7nyu | 23 | 33888 | 7yk6 | 24 | 33394 | 7xqf |

**Supplementary Table S6.**
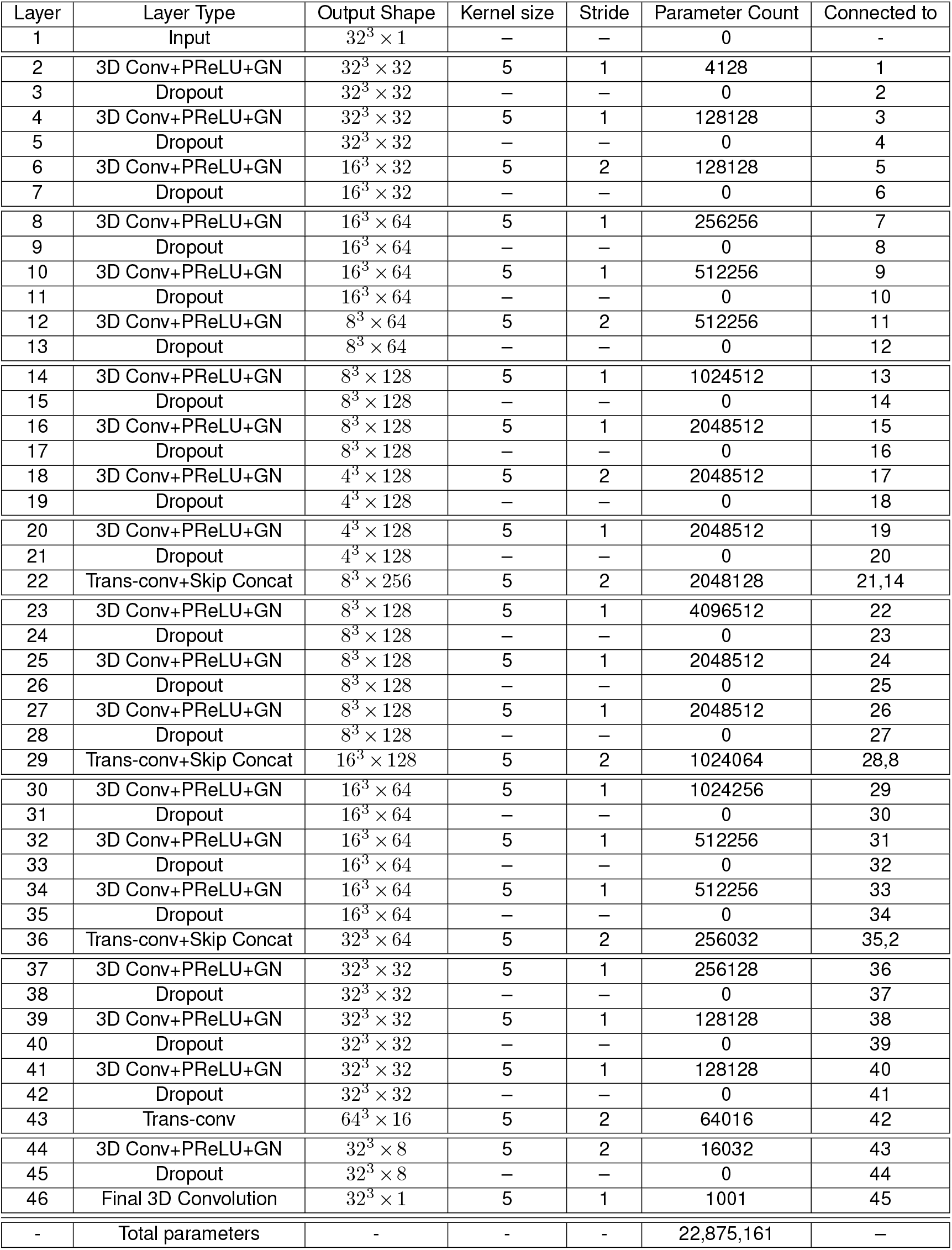
Details of MC-EMmerNet architecture.

| Layer | Layer Type | Output Shape | Kernel size | Stride | Parameter Count | Connected to |
| --- | --- | --- | --- | --- | --- | --- |
| 1 | Input | $32^3 \times 1$ | — | — | 0 | - |
| 2 | 3D Conv+PReLU+GN | $32^3 \times 32$ | 5 | 1 | 4128 | 1 |
| 3 | Dropout | $32^3 \times 32$ | — | — | 0 | 2 |
| 4 | 3D Conv+PReLU+GN | $32^3 \times 32$ | 5 | 1 | 128128 | 3 |
| 5 | Dropout | $32^3 \times 32$ | — | — | 0 | 4 |
| 6 | 3D Conv+PReLU+GN | $16^3 \times 32$ | 5 | 2 | 128128 | 5 |
| 7 | Dropout | $16^3 \times 32$ | — | — | 0 | 6 |
| 8 | 3D Conv+PReLU+GN | $16^3 \times 64$ | 5 | 1 | 256256 | 7 |
| 9 | Dropout | $16^3 \times 64$ | — | — | 0 | 8 |
| 10 | 3D Conv+PReLU+GN | $16^3 \times 64$ | 5 | 1 | 512256 | 9 |
| 11 | Dropout | $16^3 \times 64$ | — | — | 0 | 10 |
| 12 | 3D Conv+PReLU+GN | $8^3 \times 64$ | 5 | 2 | 512256 | 11 |
| 13 | Dropout | $8^3 \times 64$ | — | — | 0 | 12 |
| 14 | 3D Conv+PReLU+GN | $8^3 \times 128$ | 5 | 1 | 1024512 | 13 |
| 15 | Dropout | $8^3 \times 128$ | — | — | 0 | 14 |
| 16 | 3D Conv+PReLU+GN | $8^3 \times 128$ | 5 | 1 | 2048512 | 15 |
| 17 | Dropout | $8^3 \times 128$ | — | — | 0 | 16 |
| 18 | 3D Conv+PReLU+GN | $4^3 \times 128$ | 5 | 2 | 2048512 | 17 |
| 19 | Dropout | $4^3 \times 128$ | — | — | 0 | 18 |
| 20 | 3D Conv+PReLU+GN | $4^3 \times 128$ | 5 | 1 | 2048512 | 19 |
| 21 | Dropout | $4^3 \times 128$ | — | — | 0 | 20 |
| 22 | Trans-conv+Skip Concat | $8^3 \times 256$ | 5 | 2 | 2048128 | 21,14 |
| 23 | 3D Conv+PReLU+GN | $8^3 \times 128$ | 5 | 1 | 4096512 | 22 |
| 24 | Dropout | $8^3 \times 128$ | — | — | 0 | 23 |
| 25 | 3D Conv+PReLU+GN | $8^3 \times 128$ | 5 | 1 | 2048512 | 24 |
| 26 | Dropout | $8^3 \times 128$ | — | — | 0 | 25 |
| 27 | 3D Conv+PReLU+GN | $8^3 \times 128$ | 5 | 1 | 2048512 | 26 |
| 28 | Dropout | $8^3 \times 128$ | — | — | 0 | 27 |
| 29 | Trans-conv+Skip Concat | $16^3 \times 128$ | 5 | 2 | 1024064 | 28,8 |
| 30 | 3D Conv+PReLU+GN | $16^3 \times 64$ | 5 | 1 | 1024256 | 29 |
| 31 | Dropout | $16^3 \times 64$ | — | — | 0 | 30 |
| 32 | 3D Conv+PReLU+GN | $16^3 \times 64$ | 5 | 1 | 512256 | 31 |
| 33 | Dropout | $16^3 \times 64$ | — | — | 0 | 32 |
| 34 | 3D Conv+PReLU+GN | $16^3 \times 64$ | 5 | 1 | 512256 | 33 |
| 35 | Dropout | $16^3 \times 64$ | — | — | 0 | 34 |
| 36 | Trans-conv+Skip Concat | $32^3 \times 64$ | 5 | 2 | 256032 | 35,2 |
| 37 | 3D Conv+PReLU+GN | $32^3 \times 32$ | 5 | 1 | 256128 | 36 |
| 38 | Dropout | $32^3 \times 32$ | — | — | 0 | 37 |
| 39 | 3D Conv+PReLU+GN | $32^3 \times 32$ | 5 | 1 | 128128 | 38 |
| 40 | Dropout | $32^3 \times 32$ | — | — | 0 | 39 |
| 41 | 3D Conv+PReLU+GN | $32^3 \times 32$ | 5 | 1 | 128128 | 40 |
| 42 | Dropout | $32^3 \times 32$ | — | — | 0 | 41 |
| 43 | Trans-conv | $64^3 \times 16$ | 5 | 2 | 64016 | 42 |
| 44 | 3D Conv+PReLU+GN | $32^3 \times 8$ | 5 | 2 | 16032 | 43 |
| 45 | Dropout | $32^3 \times 8$ | — | — | 0 | 44 |
| 46 | Final 3D Convolution | $32^3 \times 1$ | 5 | 1 | 1001 | 45 |
| - | Total parameters | - | - | - | 22,875,161 | — |

